# The zinc metalloprotease ZMPSTE24 binds a distinct topological isoform of the tail-anchored protein IFITM3

**DOI:** 10.64898/2026.02.27.708584

**Authors:** Eric D. Spear, Khurts Shilagardi, Sonia Sarju, Susan Michaelis

## Abstract

The biogenesis of integral membrane proteins is complex, as revealed by an ever-growing number of cellular components shown to be dedicated to the insertion, folding, surveillance, rectification, or quality control of specific client membrane proteins. The zinc metalloprotease ZMPSTE24 and its yeast homolog Ste24 have well-established roles in the proteolytic maturation of the nuclear scaffold protein lamin A and yeast a-factor, respectively. Additionally, Ste24 has been implicated through yeast genetic screens in a variety of membrane processes, including ER- associated degradation (ERAD), Sec61 translocon “unclogging,” the unfolded protein response (UPR), and potentially as a membrane protein topology determinant. Recently, an interaction was demonstrated between ZMPSTE24 and the antiviral interferon induced transmembrane protein IFITM3, although the functional significance of this interaction is not well-understood. IFITM3 is a tail-anchored protein with a cytoplasmic N-terminus, a single transmembrane span, and a lumenal/exocellular C-terminus. Here, we show that a catalytic-dead version of ZMPSTE24, ZMPSTE24^E336A^, exhibits enhanced binding to IFITM3, and this bound species of IFITM3 is hypo-palmitoylated. Using a split fluorescence topology reporter, we demonstrate that ZMPSTE24^E336A^ “traps” and stabilizes a subpopulation of IFITM3 molecules with an atypical membrane topology, whose C-terminus is cytosolic instead of lumenal. Such inverted forms of IFITM3 are also detected in the presence of ERAD inhibitors when ZMPSTE24^E336A^ is absent. We hypothesize the ZMPSTE24^E336A^ trap mutant reveals a normally transient isoform of IFITM3 whose transmembrane span is inverted and that ZMPSTE24 is involved in the quality control of IFITM3 topology, either inverting, correcting or assisting in removal of aberrant IFITM3 molecules.

## Introduction

ZMPSTE24 is a zinc metalloprotease characterized by its seven transmembrane (TM) spans that form a large intramembrane chamber enclosing an HExxH…E active site motif (H is histidine, E is glutamate, and x is any amino acid). These residues coordinate zinc (Zn^2+^) and carry out catalysis (1–4). ZMPSTE24 is dually localized in the endoplasmic reticulum (ER) and the inner nuclear membrane and mediates the proteolytic maturation of prelamin A, the farnesylated precursor of lamin A, an important structural component of the nuclear lamina in mammals and many other vertebrates (5–7). The *Saccharomyces cerevisiae* homolog of ZMPSTE24, known as Ste24, mediates proteolytic maturation of the farnesylated precursor of the yeast mating pheromone a-factor (8–13). While yeast do not encode lamins and mammalian cells do not encode a-factor, human ZMPSTE24 and yeast Ste24 are structurally and functionally analogous, capable of cleaving both prelamin A and a-factor substrates when expressed in a yeast system (14–16).

Although prelamin A and a-factor are the only well-defined substrates for ZMPSTE24 and Ste24, several studies suggest that these integral membrane proteins may also have additional roles in protein quality control. For instance, yeast mutants lacking Ste24 (*ste24Δ*) exhibit synthetic growth defects when paired with mutations affecting other ER homeostasis pathways, including protein translocation/insertion, ER-associated degradation (ERAD), and protein folding(17, 18). A genetic screen in yeast by Tipper and Harley demonstrated that mutations in *STE24* and *SPF1* (ATP13A1 in mammals), which encodes an ER-localized ATPase, affect the topology of a single-spanning TM reporter protein (19). Consistent with functions in ER homeostasis, both *ste24Δ* and *spf1Δ* mutants exhibit chronic ER stress, as evidenced by strong activation of the unfolded protein response (UPR) under normal conditions (20). In addition, a *ste24Δ* yeast mutant or human cells treated with ZMPSTE24 inhibitors are deficient in clearance of clogged Sec61 translocons (21).

Recent studies have shown that ZMPSTE24 co-immunoprecipitates with the interferon-induced transmembrane proteins (IFITMs), which are components of the innate immune response (22–25). The IFITM proteins, including IFITM1, IFITM2, and IFITM3, modulate infection of host cells by enveloped viruses. Several studies have shown that overexpression of IFITM proteins inhibit infection, while depletion of the IFITM proteins makes cells more susceptible (26–31). However, other studies have shown that specific IFITM proteins bind viral membrane proteins and may utilize them as entry factors (31–33), indicating that the IFITMs may restrict or enhance infection, based on cell type or virus.

The IFITMs are small tail-anchored proteins ∼130 amino acids in length with a single TM span very near their C-terminus (34). The IFITMs adopt a topology with a cytosolic N-terminus and a short luminal/exocellular C-terminus (N_cyto_-C_exo_ topology) (35, 36) (see Fig. 1). Their antiviral activity depends on the presence of a short amphipathic alpha helix that binds cholesterol and stiffens membranes, thereby inhibiting fusion between viral and host cell membranes (27, 37–40). The IFITMs contain three evolutionarily conserved cysteines that are covalently modified by palmitate, a lipid modification that enhances their membrane stability and is crucial for their antiviral function (41–44). They can homo-oligomerize with themselves and one another, which may also contribute to their antiviral activity (28, 45–47). Overexpression of IFITM3 or its interactor ZMPSTE24 confers resistance against many enveloped viruses, and IFITM3 depends on ZMPSTE24 for this antiviral activity (22, 24, 25). However, the functional relationship between the IFITMs and ZMPSTE24 is poorly understood. Investigating how ZMPSTE24 influences IFITM biogenesis, activity, or membrane topology could provide insights into the essential pathways that help cells defend against viral threats.

**Figure 1.**
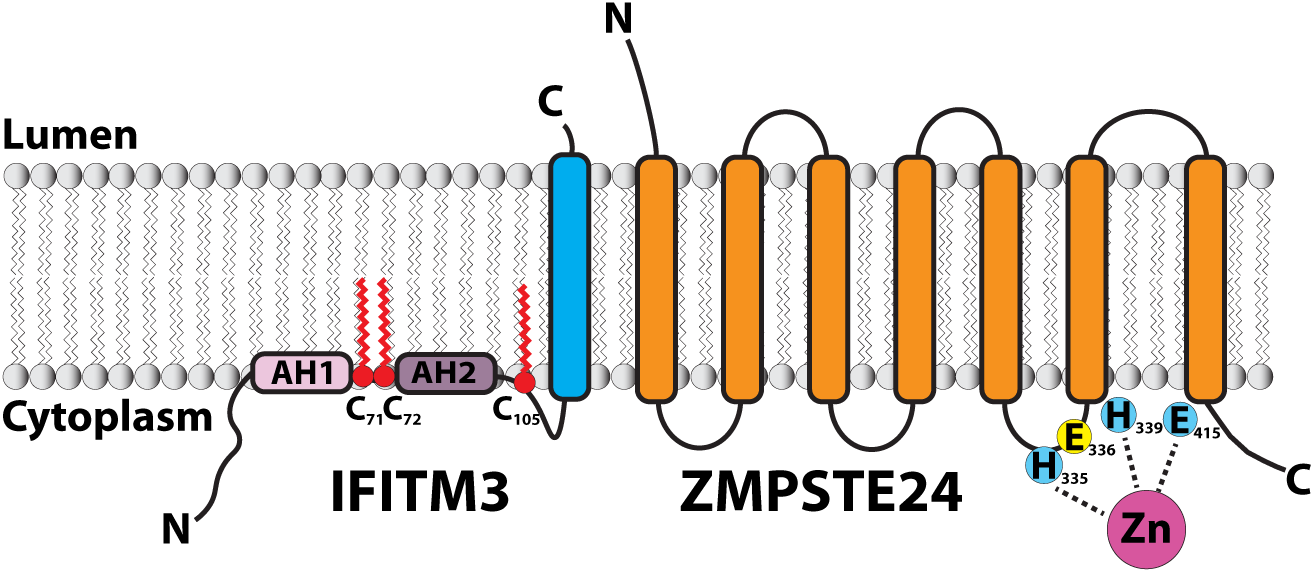
Schematic representation of IFITM3 and ZMPSTE24 in a membrane bilayer. IFITM3 has a C-terminal TM span, two N-terminal alpha-helices, AH1 (which is amphipathic) and AH2, and three conserved cysteines (C71, C72 and C105) that can undergo S-palmitoylation (red squiggly lines). ZMPSTE24 has seven TM spans and an HExxH…E zinc metalloprotease domain. The Zn^2+^ ion (pink) is coordinated by residues H335, H339 and E415 (blue). The catalytic glutamate E336 (yellow) activates a water molecule for peptide hydrolysis.

In this study, we specifically examine the interaction between ZMPSTE24 and IFITM3. Our findings confirm that the major population of IFITM3 molecules have a cytosolic N-terminus and a lumenal C-terminus (N_cyto_-C_exo_ topology) but reveal a subpopulation of IFITM3 molecules with a non-canonical topology whose C-terminus is in the cytosol (N_cyto_-C_cyto_). We show that this isoform of IFITM3 is “trapped” by a catalytically inactive mutant variant of ZMPSTE24, ZMPSTE24^E336A^, and that it is hypo-palmitoylated. Additionally, we discovered that pharmacological inhibition of enzymes involved in ubiquitin-dependent proteasomal degradation also leads to accumulation of a subpopulation of IFITM3 with a cytosolic C-terminus. Collectively, our results suggest a model in which ZMPSTE24 plays an important role in IFITM3 quality control, involved in generating, correcting or removing an IFITM3 isoform with an inverted TM span topology.

## Results

### ZMPSTE24^E336A^ efficiently ‘traps’ interacting IFITM3

A schematic of ZMPSTE24 and IFITM3 featuring their TM spans and other key features, including ZMPSTE24’s catalytic HExxH…E residues and IFITM3’s sites for palmitoylation, is shown in Fig 1. ZMPSTE24 was previously shown to co-immunoprecipitate (co-IP) with C-terminally Flag-tagged IFITM3 or untagged IFITM2 and IFITM3 induced by interferon β (22, 23, 25). We recently demonstrated that N-terminally myc-tagged IFITM3 could be immunoprecipitated with Flag-tagged ZMPSTE24, and this occurred more efficaciously using ZMPSTE24^E336A^, a catalytically inactive mutant that disrupts a glutamate (E336) in the HExxH … E active site of ZMPSTE24 (24). As indicated in Fig. 1, the active site amino acids H335, H339, and E415 (colored blue) coordinate the Zn²⁺ ion in ZMPSTE24 and E336 (colored yellow) is believed to play a direct role in positioning and activating a water molecule, which acts as a nucleophile during catalysis (1, 3, 48).

To determine whether mutations in the other residues of the HExxH … E active site co-precipitated IFITM3 comparably to ZMPSTE24^E336A^, we individually mutated H335, H339 and E415 to alanine in N-terminally Flag-tagged ZMPSTE24. Absence of proteolytic activity for each ZMPSTE24 active site mutant was confirmed using a prelamin A cleavage reporter, LMNA_431-664_ (Fig. S1A). Co-IP of myc-tagged IFITM3 by these Flag-tagged ZMPSTE24 mutants reveals that in comparison to the ZMPSTE24^E336A^ mutant, neither wild-type ZMPSTE24 nor the other active site mutants showed enhanced IFITM3 binding, displaying ∼3-13% of the level of interaction by co-IP as compared to ZMPSTE24^E336A^-IFITM3 (Fig. 2A; compare lane 4 to lanes 2, 3, 5, and 6). Notably, a faster migrating myc-IFITM3 band is observed in the lysate only when co-expressing ZMPSTE24^E336A^ (Fig. 2A, lane 4, arrow), suggesting this species is the form co-IP’d with ZMPSTE24 ^E336A^. Introduction of the H339A or E415A mutations into ZMPSTE24^E336A^ ablated its enhanced IFITM3 binding (Fig. 2A, compare lanes 7 and 8 to lane 4); thus, zinc coordination, and not absence of catalytic activity per se, may be required for IFITM3 “trapping” by ZMPSTE24^E336A^. In addition, Flag-tagged ZMPSTE24^E336A^ also co-IP’d endogenous and ectopically-expressed IFITM1, 2, and 3 more efficaciously than wild-type ZMPSTE24 (Fig. S1B and C), although the difference was modest in the case of IFITM1. Finally, Flag-tagged IFITM3 co-IP’d myc-ZMPSTE24^E336A^ more efficiently than wild-type myc-ZMPSTE24, confirming the specificity of the interaction in both directions (Fig. S1D, compare lane 3 to lane 2). Taken together, these studies indicate that the ZMPSTE24^E336A^ mutant “traps” the interaction between ZMPSTE24 and IFITM3 with high efficiency. Therefore, we used this version of ZMPSTE24 for the studies below.

**Figure 2.**
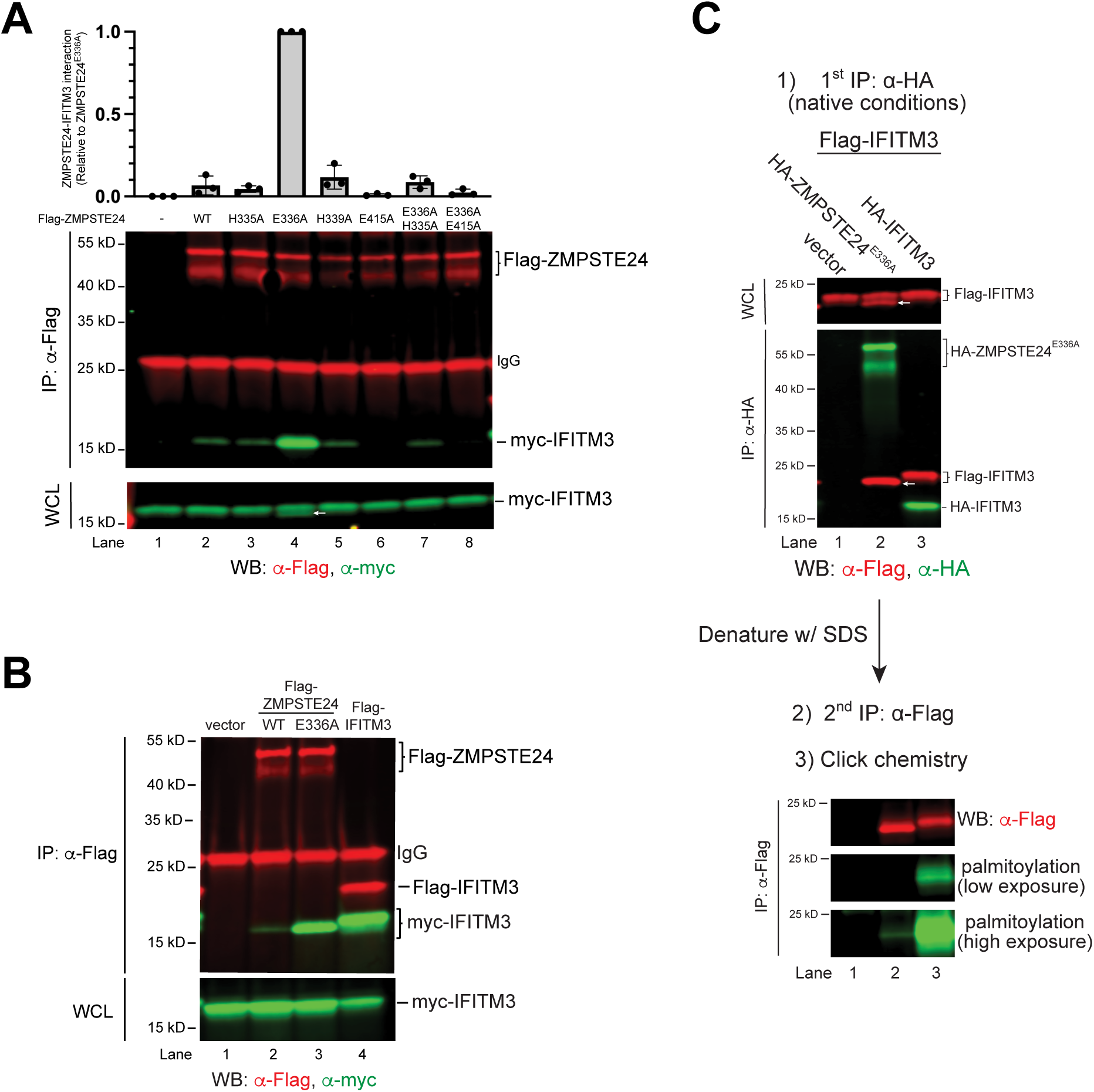
The mutant protein ZMPSTE24^E336A^, but not other HExxH…E mutants, shows enhanced binding to IFITM3 and the bound species is hypo-palmitoylated. (A) The ZMPSTE24^E336A^ mutant co-IPs IFITM3 more effectively than WT ZMPSTE24 or other active site mutants. Vector or the indicated Flag-tagged ZMPSTE24 variants were co-transfected with myc-tagged IFITM3 in HEK293 cells and subjected to co-IP with anti-Flag agarose, SDS-PAGE, and Western blotting with anti-Flag and anti-myc antibodies. Flag-ZMPSTE24 (red) and myc-IFITM3 (green) proteins are indicated in the IP and whole cell lysate (WCL). The white arrow (Lane 4) indicates faster migrating myc-IFITM3 species only observed when co-expressing ZMPSTE24^E336A^. The ZMPSTE24-IFITM3 interaction was quantified from three independent experiments, with the average and standard deviation of the mean shown (normalized to ZMPSTE24^E336A^). It should be noted that ZMPSTE24, whether tagged, or untagged always migrates as 2 bands. (B) The myc-IFITM3 species that co-IP’s with Flag-ZMPSTE24 ^E336A^ migrates more rapidly than the species which co-IP’s with Flag-IFITM3. Vector, Flag-ZMPSTE24, Flag-ZMPSTE24^E336A^, or Flag-IFITM3 were co-transfected with myc-tagged IFITM3 in HEK293 cells. Proteins were analyzed as described in (A). It should be noted that this panel corresponds to lanes 9-12 in Fig. S1C. (C) ZMPSTE24^E336A^ binds a hypo-palmitoylated form of IFITM3 compared to the form of IFITM3 in homo-oligomers. Cells expressing vector, HA-tagged ZMPSTE24^E336A^ or HA-IFITM3 together with Flag-IFITM3 were metabolically labeled with 17-ODYA prior to native co-IP (to maintain the complex), using anti-HA agarose beads. A portion of the immune complexes were analyzed by Western blotting with anti-Flag and anti-HA antibodies (top). White arrow (lane 2, WCL and IP) indicates faster migrating Flag-IFITM3 species only observed when co-expressing HA-ZMPSTE24^E336A^. IP’d proteins were denatured with SDS, re-IP’d with anti-Flag agarose, and subjected to copper-mediated click chemistry with an azide-linked IRDye800 fluorophore followed by SDS-PAGE and Western blotting (bottom). A palmitoylation fluorescent signal for IFITM3 co-IP’d with HA-IFITM3 is evident, but a very faint signal for IFITM3 co-IP’d with ZMPSTE24^E336A^ can only be observed upon increased exposure (compare lanes 2 and 3).

### ZMPSTE24^E336A^ interacts with a hypo-palmitoylated form of IFITM3

IFITMs are known to form both homo- and hetero-oligomers with other IFITMs. In our study, we noticed that the myc-IFITM3 species that co-IP’s with Flag-ZMPSTE24^E336A^ resolves as a single, faster migrating band than the myc-IFITM3 species that co-IP’s with Flag-IFITM3 in the homo-oligomer (Fig. 2B, compare lanes 3 and 4). A similar pattern was observed for IFITM1 and IFITM2 (Fig. S1C).

Given that IFITM3 can be palmitoylated on one, two, or all three of its conserved cysteines (42, 44), we hypothesized that the faster migrating species of IFITM3 co-precipitated by ZMPSTE24^E336A^ may be under-palmitoylated compared to IFITM3 in the homo-oligomer. To compare the palmitoylation status of the IFITM3 species co-precipitated by ZMPSTE24^E336A^ versus IFITM3, we performed click-chemistry (49, 50) (Fig. 2C). Cells expressing the indicated proteins were metabolically labeled with an alkyne-functionalized fatty acid palmitate analog, 17-octadecanoic acid (17-ODYA), as described in the Experimental procedures. As expected, both slower and faster-migrating species of Flag-IFITM3 were observed in the whole cell lysate co-expressing HA-ZMPSTE24^E336A^ (Fig. 2C, top panel, lane 2), and the Flag-IFITM3 species co-IP’d by HA-ZMPSTE24^E336A^ migrates slightly faster than that co-purified by HA-IFITM3 (Fig. 2C, top panel, compare lanes 2 and 3). To specifically analyze the palmitoylation status of Flag-IFITM3 in the immune complexes, immunoprecipitated proteins were denatured with SDS, and then re-immunoprecipitated with anti-Flag beads, followed by copper-mediated click chemistry with an azide-linked fluorophore (IRDye800 CW azide) and analysis by SDS-PAGE (Fig. 2C, bottom panel). Despite similar levels of Flag-IFITM3 in both samples, we found that HA-ZMPSTE24^E336A^ pulled down a hypo-palmitoylated species compared to the Flag-IFITM3 species in the homo-oligomer (Fig. 2C, bottom panel, compare lanes 2 and 3).

### A cysteine crosslink between ZMPSTE24^E336A^ and IFITM3 suggests an unconventional membrane topology for IFITM3

We observed an unexpected high molecular weight band in our co-IPs between Flag-ZMPSTE24^E336A^ and myc-IFITM3 when samples were resolved under non-reducing conditions; however, this band was absent under reducing conditions (Fig. 3A). The molecular weight (∼65 kDa) and presence of both ZMPSTE24^E336A^ and IFITM3 (Fig. 3A, lane 2, merge) suggest that the species represents an intermolecular disulfide-linked complex between the two proteins (Flag-ZMPSTE24^E336A^ migrates ∼45-50 kDa and myc-IFITM3 migrates ∼17 kDa). This complex likely forms *in vitro*, since it is primarily observed when lysis buffer includes Triton X-100, a detergent known to contain small amounts of peroxides, which are oxidizing agents (51). Notably, the complex is not detected when higher-purity detergents, such as dodecyl maltoside and digitonin are used in the lysis buffer (see Fig. S2). Furthermore, the presence of N-ethylmaleimide (NEM) in the lysis buffer, which modifies free thiols and inhibits cysteine oxidation, effectively prevents the formation of the intermolecular disulfide complex (Fig. 3B). Together these data support the identity of the higher molecular weight band as an intermolecular disulfide complex between ZMPSTE24^E336A^ and IFITM3.

**Figure 3.**
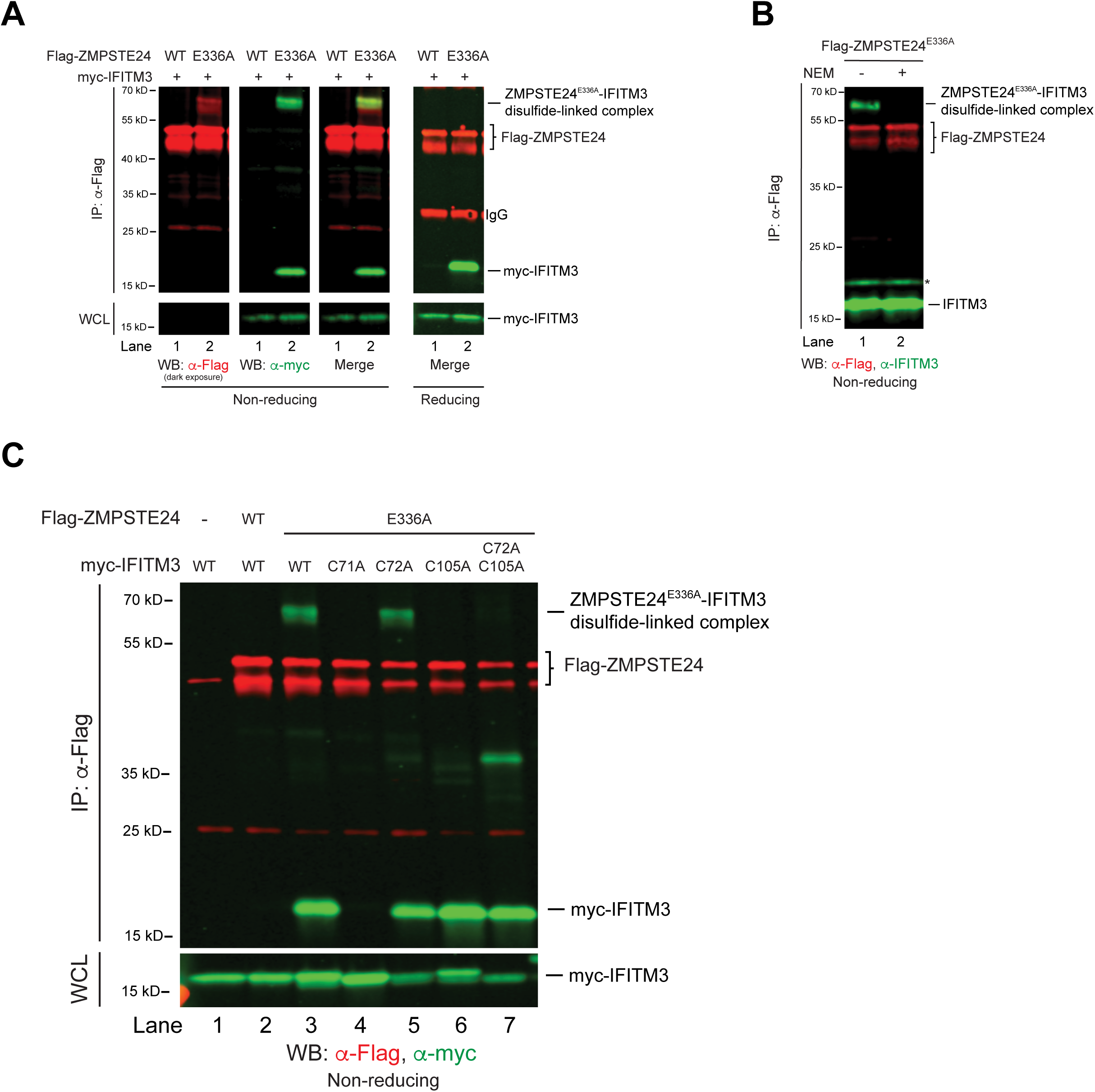
IFITM3 Cys105 forms an intermolecular disulfide bond with ZMPSTE24^E336A^ *in vitro.* (A) HEK293 cells transfected with Flag-ZMPSTE24 or Flag-ZMPSTE24^E336A^ and myc-IFITM3 were subjected to co-IP with anti-Flag agarose beads. Immune complexes were resolved by non-reducing (left) or reducing (right) SDS-PAGE prior to Western blotting with anti-Flag and anti-myc antibodies. A higher molecular weight species (∼65 kDa) containing both the myc and Flag signals is detected exclusively under non-reducing conditions. (B) The disulfide-linked species containing ZMPSTE24^E336A^ and IFITM3 is disrupted by NEM in the lysis buffer. Cells transfected with Flag-ZMPSTE24^E336A^ were induced with interferon β for 18 hrs prior to lysis with 1% Triton X-100 buffer without (-) or with (+) 0.1mM NEM. Proteins were immunoprecipitated with anti-Flag agarose beads, resolved by non-reducing SDS-PAGE and analyzed by Western blotting with anti-Flag and anti-IFITM3 antibodies. * denotes a non-specific background band. (C) HEK293 cells were transfected with vector, Flag-ZMPSTE24 or Flag-ZMPSTE24^E336A^ with the indicated myc-tagged IFITM3 cysteine variants and subjected to co-IP followed by non-reducing SDS-PAGE and Western blotting. The IFITM3-C71A mutant fails to bind ZMPSTE24 altogether, and the ZMPSTE24^E336A^-IFITM3 intermolecular disulfide complex is absent for the IFITM3-C105A mutant.

Although this disulfide-linked species may only form *in vitro*, we hypothesized that identifying the cysteines in IFITM3 and ZMPSTE24^E336A^ that are close enough to oxidize (within ∼5Å), could yield insights into the interaction between IFITM3 and ZMPSTE24^E336A^. Furthermore, our finding that the form of IFITM3 that co-precipitates with ZMPSTE24^E336A^ is hypo-palmitoylated supports the notion that at least one of the conserved cysteines in IFITM3 may be unmodified and thus available to disulfide bond with ZMPSTE24. To determine which cysteine in IFITM3 oxidized with ZMPSTE24^E336A^, we mutated each of the three cysteines in myc-IFITM3 to alanine and expressed these variants with Flag-ZMPSTE24^E336A^ for co-IP experiments. IFITM3-C71A failed to co-IP with ZMPSTE24^E336A^ altogether (Fig. 3C compare lanes 3 and 4), so it was not relevant for this analysis and is discussed further below. Whereas IFITM3-C72A and IFITM3-C105A were efficiently co-IP’d by Flag-ZMPSTE24^E336A^, the high molecular weight intermolecular disulfide complex was present for IFITM3-C72A but absent for IFITM3-C105A and the double mutant IFITM3-C72A, C105A (Fig. 3C, compare lanes 5-7). This result indicates it is Cys105 in IFITM3 that is oxidized to ZMPSTE24^E336A^.

We next turned our attention to ZMPSTE24. Although human ZMPSTE24 has five cysteines (C109, C176, C324, C359 and C406; Fig. 4A), none are highly conserved among ZMPSTE24 homologs. Importantly, a version of ZMPSTE24^E336A^ devoid of cysteines (Cys-less) co-precipitated IFITM3 to a similar extent as WT ZMPSTE24^E336A^ (Fig. 4B, compare lanes 7 and 1), indicating disulfide bonding is not required for the interaction between ZMPSTE24 and IFITM3.

**Figure 4.**
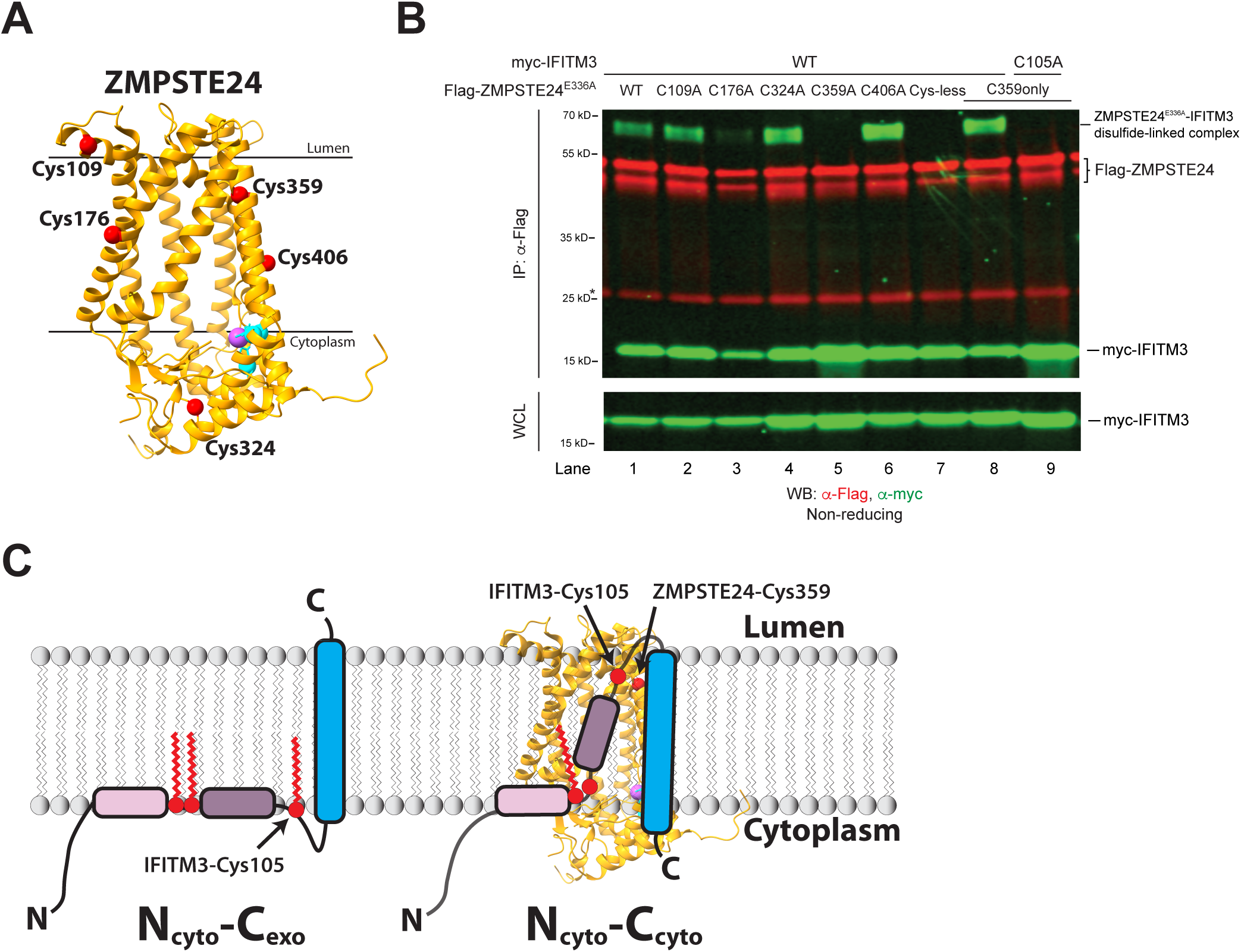
ZMPSTE24-Cys359 is necessary and sufficient to form the intermolecular disulfide bond with IFITM3-Cys105. (A) X-ray crystal structure (pdb: 5SYT) (1) of ZMPSTE24. Cysteines (red balls), Zn^2+^ (purple ball), and Zn^2+^-coordinating residues H335, H339 and E415 (blue) are indicated. Approximate position of membranes separating cytoplasm and lumen is shown. (B) Cys359 in ZMPSTE24^E336A^ is required for disulfide bonding to IFITM3. The indicated Flag-tagged ZMPSTE24 ^E336A^ Cys mutants were co-expressed with myc-IFITM3 (or myc-IFITM3^C105A^, Lane 9) and subjected to co-IP and non-reducing SDS-PAGE as described above. ZMPSTE24^E336A^ lacking cysteines (“Cys-less”) and C359A bind IFITM3 but do not form the intermolecular disulfide species. (C) Working model for the orientation of IFITM3’s TM span in the absence or presence of ZMPSTE24^E336A^. Schematic of fully palmitoylated IFITM3 in the N_cyto_-C_exo_ topology (left) and hypo-palmitoylated IFITM3 in the N_cyto_-C_cyto_ topology bound to ZMPSTE24 (right). Positions of the cysteines in IFITM3 and ZMPSTE24^E336A^ that oxidize during co-IP are indicated.

To determine which cysteine in ZMPSTE24^E336A^ forms the disulfide bond with IFITM3-Cys105, we mutated individual cysteines to alanine in Flag-ZMPSTE24^E336A^ and used these mutants to co-IP myc-IFITM3 under non-reducing conditions. All of the Cys substitution mutants precipitated the ∼17kD myc-IFITM3 species (Fig. 4B, lanes 2-6), but only the ZMPSTE24^E336A^-C359A mutant (and the cysteine-less mutant) failed to exhibit the high molecular weight disulfide-linked species with IFITM3 (Fig. 4B, lanes 5 and 7). Although the ZMPSTE24^E336A^-C176A mutant appears to be unstable, with less ZMPSTE24 present, a faint disulfide-linked species with IFITM3 can still be observed (Fig. 4B, lane 3). Furthermore, a ZMPSTE24^E336A^ mutant with only a single cysteine (C359 only) preserved the intermolecular disulfide species with IFITM3 (Fig. 4B, lane 8) and this was disrupted when using IFITM3-C105A (Fig. 4B, lane 9). Together, these data indicate that Cys359 in ZMPSTE24^E336A^ is both necessary and sufficient to form the intermolecular disulfide bond with IFITM3 Cys105.

Given the known structure of ZMPSTE24, our results have topological implications for the IFITM3 that is bound to it. Although multiple topologies have been suggested for IFITM3, recent analysis strongly indicates it conforms to the expected orientation for a typical tail-anchored protein with its N-terminus in the cytoplasm and its C-terminus in the luminal/exocellular space (an N_cyto_-C_exo_ orientation) (Fig. 4C, left) (35, 36). Our findings indicate that Cys359 in ZMPSTE24^E336A^, which is located in or near the ER lumen, can oxidize with Cys105 in IFITM3, a cysteine predicted to be cytoplasmic. This suggests an unexpected “flipped” topology for the TM span of IFITM3 (N_cyto_-C_cyto_) when it is bound to ZMPSTE24^E336A^ (see Fig. 4C, right).

### Topology analysis of IFITM3 in the presence of ZMPSTE24^E336A^ reveals a subpopulation whose C-terminus is cytosolic

To directly examine the topology of IFITM3 in the presence and absence of ZMPSTE24^E336A^, we utilized a split-fluorescence topology reporter system devised by McKenna et al. (52) for analyzing the orientation of proteins with a single TM span (Fig 5A). In this reporter system, HEK293 cells stably express the first 10 ²-strands of mNeon Green3K (referred to here as GFP) in the cytoplasm and the first 10 ²-strands of mCherry in the ER lumen (Fig. 5A, left). The test protein, IFITM3 in our case, is tagged with tandem GFP-²11 and mCherry-²11 strands at either its N-terminus (N-²11-IFITM3) or C-terminus (IFITM3-²11-C) (Fig. 5A, right), such that fluorescence occurs when the β11 strands bind the GFP(1–10) in the cytoplasm or mCherry(1–10) in the ER lumen (Fig. 5A, left). Co-expression of BFP in these constructs, separated from the *IFITM3* genes by a P2A ribosome skipping sequence (Fig. 5A, right), allows for quantification of the fluorescent signals by flow cytometry analysis.

**Figure 5.**
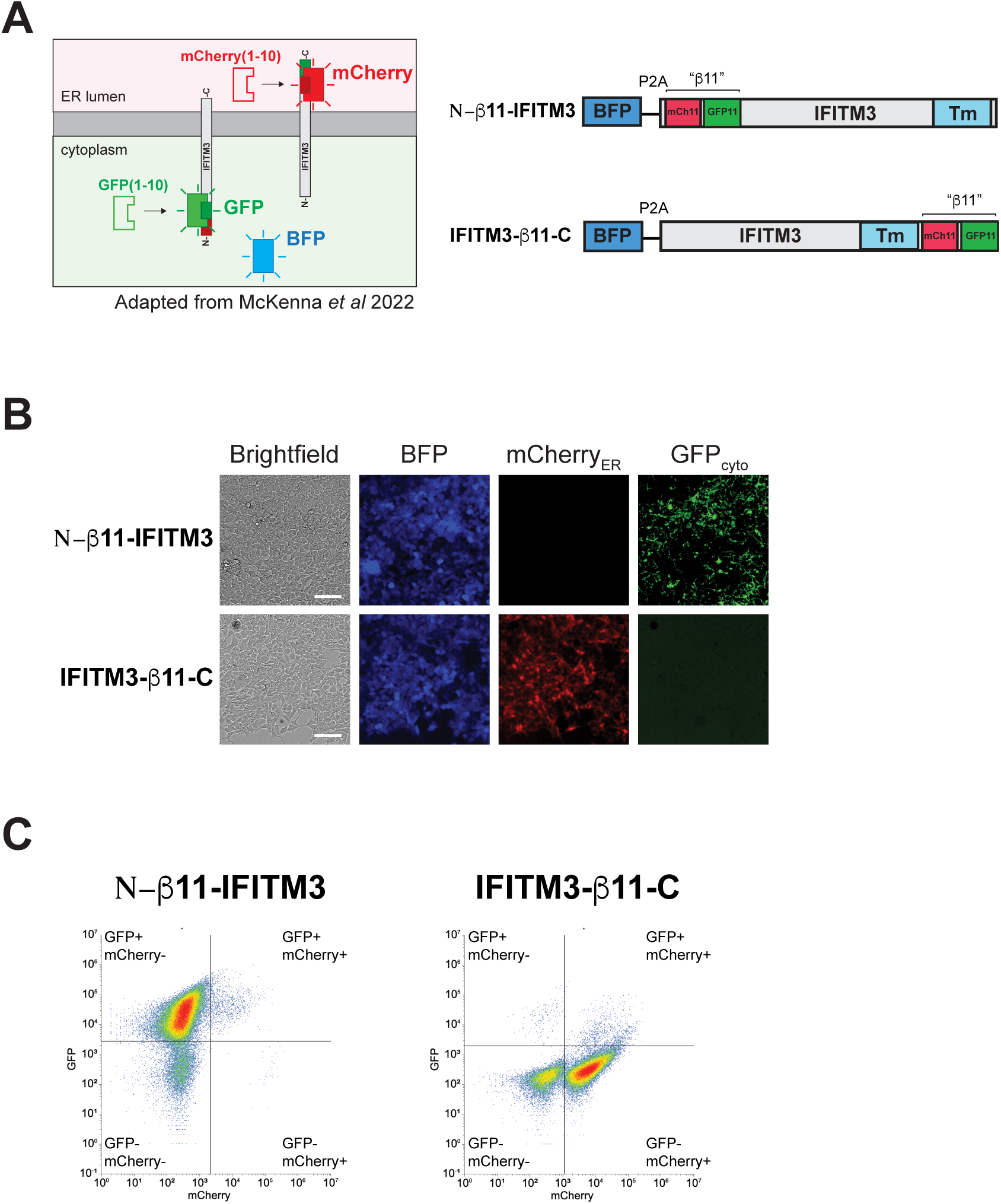
A split fluorescence topology reporter corroborates the expected N_cyto_-C_exo_ topology for IFITM3. (A) Schematic of the split fluorescence reporter system developed by Mckenna et al. (52). To determine the cellular location of the N- and C-termini of IFITM3, the indicated plasmids expressing BFP and IFITM3 tagged with the β11 strands of mCherry and GFP at the N- or C-termini were stably transfected into 293 Flp-In TRex cells expressing cytosolic GFP_1-10_ and ER-localized mCherry_1-10_. Only when the β11 peptide binds to its partner protein does fluorescence occur, indicating cytoplasmic (GFP+) or ER lumenal (mCherry+) localization. (B) Fluorescence microscopy and (C) flow cytometry of cells expressing N-β11-IFITM3 or IFITM3-β11-C proteins of a representative experiment confirm an N_cyto_-C_exo_ topology under steady state conditions. Scale bar = 100 μm. In C, the fluorescence patterns of cells are indicated for each quadrant.

We first validated the published steady-state topology of IFITM3 by fluorescence microscopy and flow cytometry (Figs. 5B and C) in wild-type cells. Cells expressing N-²11-IFITM3 displayed a predominantly GFP signal (Fig. 5B, top), while cells expressing IFITM3-²11-C displayed a predominantly mCherry signal (Fig. 5B, bottom), indicating the expected N_cyto_-C_exo_ topology for the tail anchor protein IFITM3. Flow cytometry reflected the patterns observed by microscopy for both constructs (Fig. 5C).

We next tested whether the orientation of IFITM3 would be altered by ZMPSTE24^E336A^ co-expression. Cells expressing N-²11-IFITM3 were primarily GFP-positive and mCherry negative when co-expressing ZMPSTE24^E336A^, indicating the cytoplasmic N-terminus is not altered when ZMPSTE24^E336A^ is present (Fig. 6A and D). In contrast, co-expression of ZMPSTE24^E336A^ with IFITM3-²11-C resulted in the striking appearance of a significant population of GFP-positive cells by fluorescence microscopy (Fig. 6B). Quantification of the GFP signal by flow cytometry showed that the mean GFP:BFP ratio increased ∼6-13-fold in cells expressing ZMPSTE24^E336A^, indicating an increased number of cells in which the IFITM3 C-terminus is cytoplasmically oriented. ZMPSTE24^E336A^ “trapping” of IFITM3 was required to observe this high C_cyto_ topology signal, since only a very weak effect on the topology of IFITM3-β11-C was detected in cells transfected with wild-type ZMPSTE24 (Fig. 6D and Fig. S3). We also examined the topology of IFITM3^C71A^, the mutant form of IFITM3 that we observed to poorly co-IP with ZMPSTE24^E336A^ (Fig. 3C, lane 4). In contrast to IFITM3-²11-C, the topology of IFITM3^C71A^-²11-C was unaffected by expression of ZMPSTE24^E336A^ (Fig. 6C, D and Fig. S3), providing further support that interaction and trapping of IFITM3 by ZMPSTE24^E336A^ is required to observe the “flipped” N_cyto_-C_cyto_ topology (shown in Fig.4C, right).

**Figure 6.**
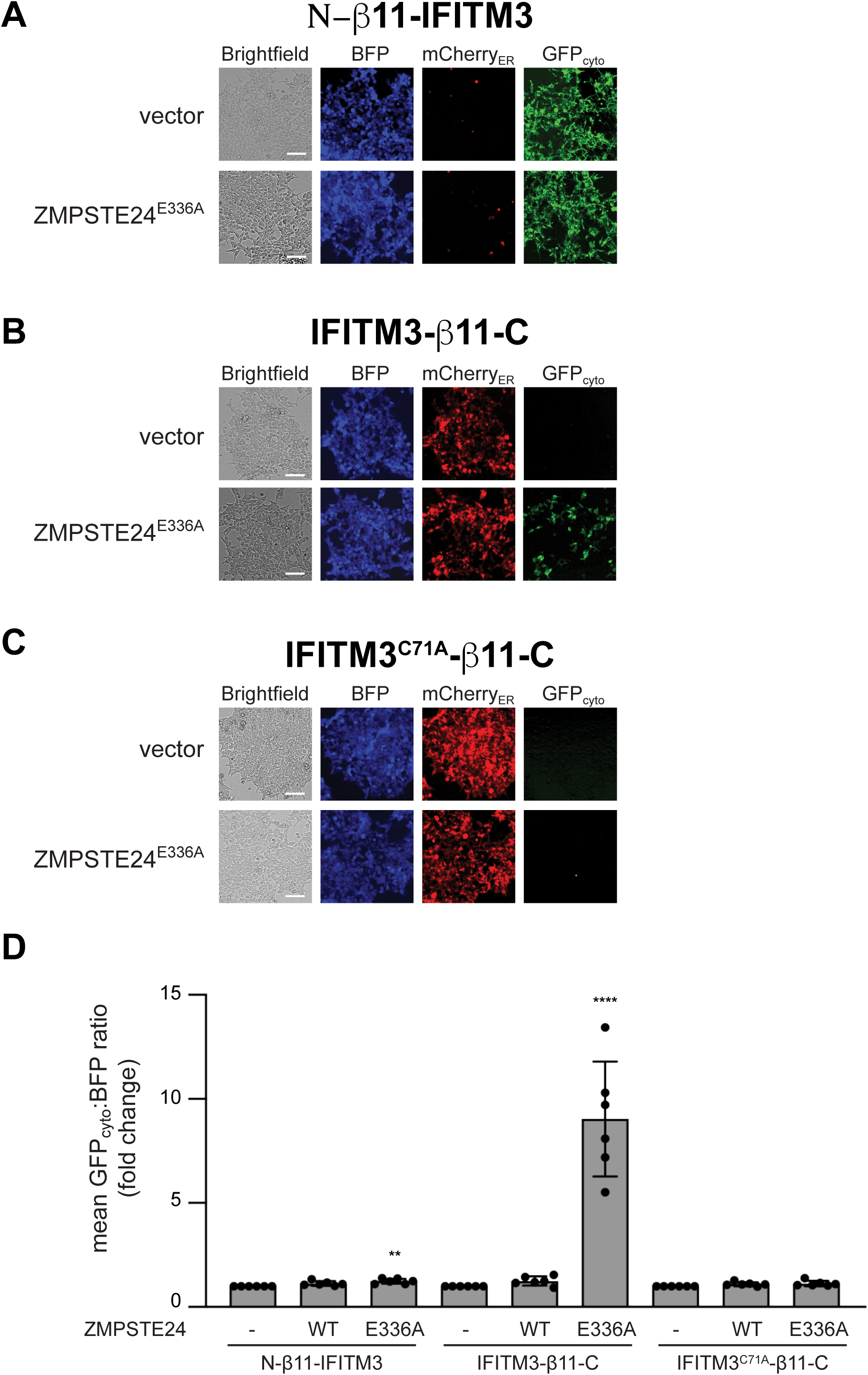
The C-terminus of IFITM3 is cytoplasmic in ZMPSTE24^E336A^-expressing cells suggesting an inverted orientation for IFITM3’s TM span. (A-C) Topology of IFITM3 constructs in the absence (vector) or presence of ZMPSTE24^E336A^. Flp-In 293 TREx topology reporter cells expressing (A) N-β11-IFITM3, (B) IFITM3-β11-C or (C) IFITM3^C71A^-β11-C were transfected with vector control or ZMPSTE24^E336A^ and analyzed by fluorescence microscopy. Scale bar = 100 μm. (D) Graph of cells in A-C were analyzed by flow cytometry (shown in Fig. S3). The mean GFP:BFP ratio was calculated for each condition, and the average and standard deviation fold change (n=6) compared to vector alone, set to 1, is shown. ** p<0.005, **** p<0.0005 in comparison to vector control (-).

### The C-terminus of IFITM3 molecules that interact with ZMPSTE24^E336A^ enter the ER lumen, presumably prior to localization in the cytosol

We envisioned two potential explanations for the observed N_cyto_-C_cyto_ topology of a subset of IFITM3 molecules in the presence of ZMPSTE24^E336A^. First, IFITM3 may initially be inserted into the ER with an N_cyto_-C_exo_ orientation, which is typical for tail-anchored proteins. Subsequently, its TM region may flip to form the N_cyto_-C_cyto_ isoform. In the latter scenario, ZMPSTE24^E336A^ might play a role in facilitating and/or stabilizing this flipped state of IFITM3. Alternatively, it is possible that the subset of IFITM3 bound to ZMPSTE24^E336A^ was incorrectly inserted into the ER membrane with an N_cyto_-C_cyto_ orientation, preventing the C-terminus from being exposed to the ER lumen. To distinguish between these possibilities, we placed an opsin glycosylation tag at the C-terminus of myc-IFITM3 (Fig. 7A). N-linked glycosylation of the opsin tag on IFITM3 (detectable as decreased gel mobility by SDS-PAGE) would indicate that the IFITM3 C-terminus had inserted into the ER lumen. A construct with the opsin tag near the N-terminus was included as a negative control, as it should never be exposed to the ER lumen.

**Figure 7.**
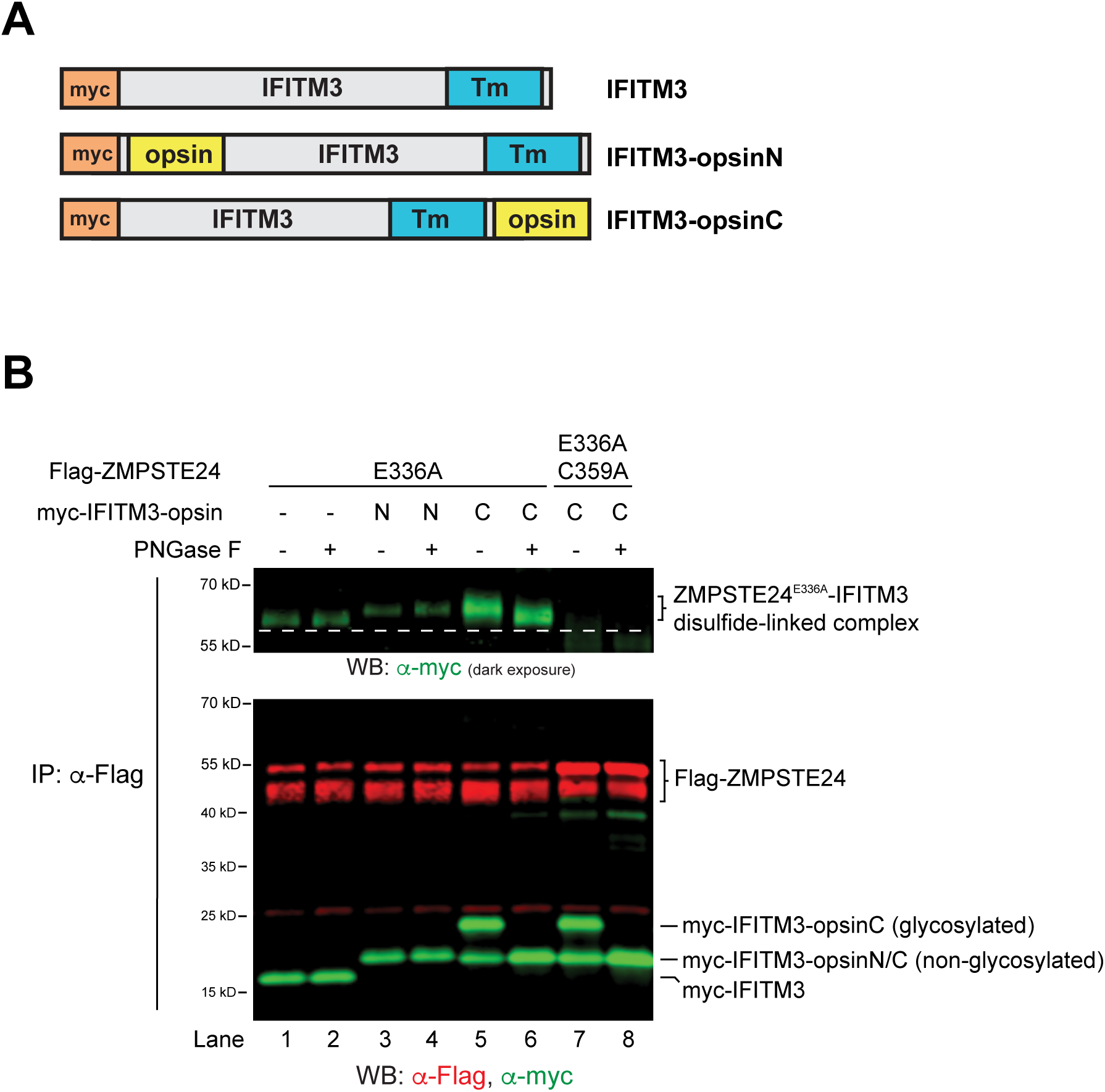
The C-terminus of IFITM3 bound to ZMPSTE24^E336A^ was exposed to the ER lumen during its biogenesis. (A) Schematic of myc-tagged IFITM3 proteins with no opsin glycosylation tag, or an opsin tag at the N- or C-terminus. (B) Co-IP by Flag-tagged ZMPSTE24^E336A^ of myc-IFITM3 proteins with N- and C-terminal opsin tags. Immune complexes were mock-treated (-) or treated with PNGase F (+) prior to non-reducing SDS-PAGE and Western blotting. The opsin tag at the C-terminus of IFITM3 is glycosylated and shifts upon PNGase treatment, as is also the case for the ∼65kDa intermolecular disulfide complex (absent when using ZMPSTE24^E336A, C359A^, lanes 7 and 8). A dotted line was added to highlight protein gel mobility differences.

Cells co-expressing Flag-ZMPSTE24^E336A^ and opsin-tagged myc-IFITM3 proteins were subjected to anti-Flag immunoprecipitation, and the immune complexes were either mock-treated or treated prior to SDS-PAGE with PNGase F, a glycosidase that removes N-linked glycosylation. As expected, the N-terminal opsin-tagged myc-IFITM3 was slightly larger with a slower gel mobility than myc-IFITM3, due to the additional tag residues, but was not glycosylated (Fig. 7B, compare lanes 3 and 4 to lanes 1 and 2, bottom panel). In contrast, roughly half of the C-terminally opsin-tagged myc-IFITM3 that co-IPs with Flag-ZMPSTE24^E336A^ is a higher molecular weight species that is sensitive to PNGase F, indicating its C-terminus had entered the ER lumen and was glycosylated (Fig. 7B, bottom panel, compare lanes 5 and 6 to lanes 3 and 4). This glycosylated species was also sensitive to Endo H, indicating it remained in the ER membrane or became inaccessible to Golgi-modifying enzymes that render glycans Endo H-resistant (Fig. S4). Furthermore, the high molecular weight ZMPSTE24^E336A^-IFITM3 disulfide-linked complex (representing the N_cyto_-C_cyto_ isoform) also shifted in molecular weight upon PNGase F treatment (Fig. 7B top panel, compare lanes 5 and 6), suggesting IFITM3 in this complex had its C-terminus exposed to the ER lumen at some point during its biosynthesis.

Taken together, the results shown in Fig. 6 and Fig. 7 favor a model whereby IFITM3 is initially inserted into the ER membrane as expected for a prototypical tail-anchored protein (N_cyto_-C_exo_) (see Fig. 4C, left). After membrane insertion, it appears that a subpopulation of IFITM3 can undergo TM span inversion, creating the N_cyto_-C_cyto_ isoform of IFITM3. We cannot distinguish at this point whether ZMPSTE24 is directly involved in the inversion of IFITM3’s TM span and/or whether ZMPSTE24^E336A^ traps and stabilizes the inverted form of IFITM3 (see Fig. 4C, right).

### ERAD Inhibitors allow detection of a topologically flipped form of IFITM3 in the absence of ZMPSTE24^E336A^

We considered the possibility that the N_cyto_-C_cyto_ inverted topological isoform of IFITM3 observed only in the presence of ZMPSTE24^E336A^ might occur in its absence, but could be unstable, and thus transient, due to ubiquitin-proteasome mediated ERAD. We therefore used the topology reporter system to test whether pharmacological inhibitors of ubiquitination, ER extraction, or proteasome activity might stabilize and allow detection of the N_cyto_-C_cyto_ IFITM3 isoform in the absence of ZMPSTE24^E336A^.

Cells expressing IFITM3-β11-C were treated for 4 hours with vehicle (DMSO) or pharmacological inhibitors of the E1 ubiquitin-activating enzyme (MLN4273), the AAA+ ATPase VCP/p97 (NMS-873), or the proteasome (bortezomib) and analyzed by flow cytometry. For each drug treatment, the mean GFP:BFP ratio increased ∼4-fold (Fig. 8A and B and Fig. S5), indicating accumulation of the IFITM3 C-terminus in the cytoplasm. Accumulation of this C_cyto_ isoform of IFITM3 was dependent on new protein synthesis, as cycloheximide treatment blocked its appearance by the p97/VCP inhibitor (Fig. 8A), suggesting that TM inversion and degradation occur soon after IFITM3 biosynthesis. Importantly, only the C-terminus of IFITM3 was strongly affected by drug treatment, since N-²11-IFITM3 and ASGR1-β11-C, a type II (non-tail-anchored) TM protein with a longer luminal/exocellular C-terminus (also N_cyto_-C_exo_), were only modestly affected (Fig. 8B and Fig. S5). Taken together, our findings suggest that the N_cyto_-C_cyto_ topological isoform of IFITM3 occurs under normal conditions but is transient because it is targeted for ubiquitin-mediated proteasome degradation.

**Figure 8.**
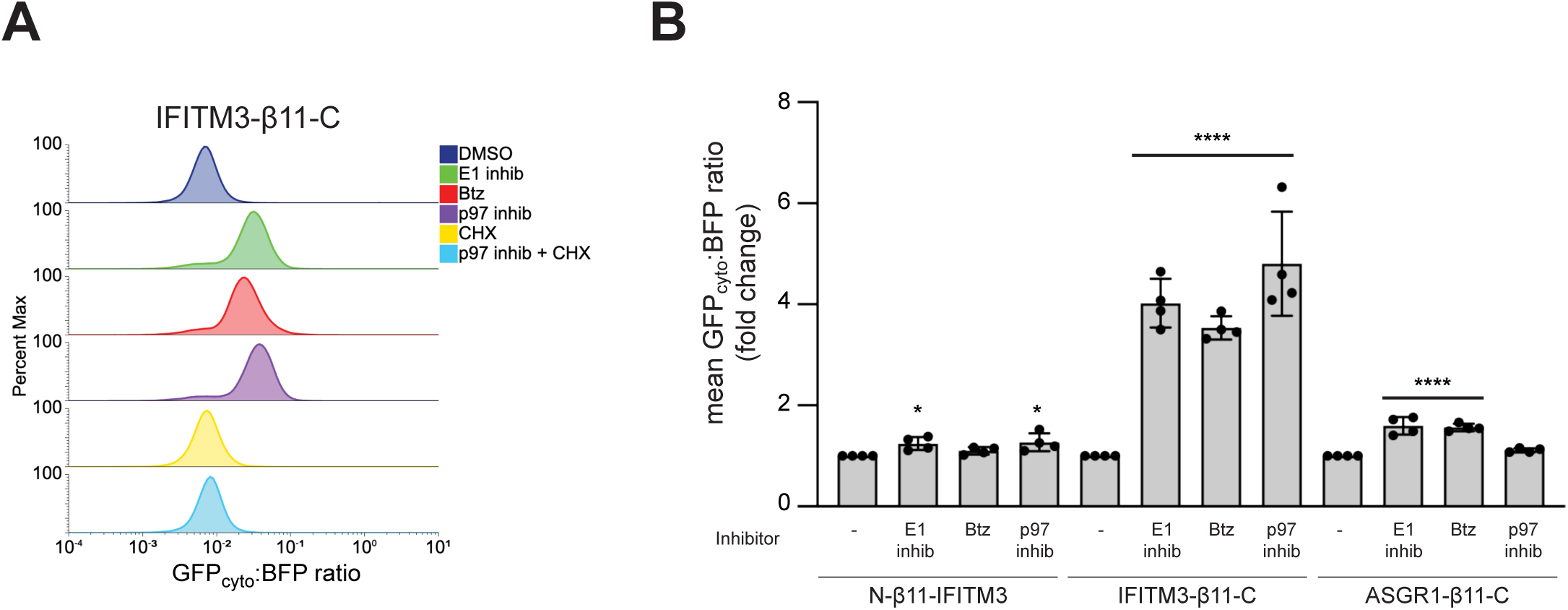
The IFITM3 N_cyto-_C_cyto_ topological isoform is unstable and degraded in a ubiquitin/proteasome-dependent manner. **(A)** Flow cytometry histogram of Flp-In 293 TREx cells expressing IFITM3-β11-C analyzed after 4 hrs of treatment with vehicle (DMSO) or inhibitors of the E1 ubiquitin-activating enzyme (E1 inhib), proteasome (Btz), p97/VCP (p97 inhib), or the protein synthesis inhibitor cycloheximide (CXH) in the absence or presence of p97 inhibitor. The GFP:BFP ratio increases upon inhibition of the ubiquitin-proteasome system. (B) Cells expressing N-β11-IFITM3, IFITM3-β11-C, or ASGR1-β11-C were treated with the indicated drugs for 4 hrs prior to analysis by flow cytometry. The average and standard deviation of the fold change GFP:BFP ratio over DMSO vehicle is shown (n=4). * p<0.05, **** p<0.0005 in comparison to DMSO treatment (-).

## Discussion

Tail anchored proteins destined for the secretory pathway, such as IFITM3, have a single hydrophobic TM span followed by a short hydrophilic segment at their C-terminus. They are inserted into the ER membrane by specialized pathways, either the GET or EMC, with their C-terminus in the luminal/exocellular space and their N-terminus in the cytosol (N_cyto_-C_exo_) (reviewed in (53, 54)). Here we validate that IFITM3 is predominantly in the expected N_cyto_-C_exo_ orientation. However, importantly, we demonstrate that a subpopulation of IFITM3 exists in which IFITM3’s TM span has acquired the reverse orientation after membrane insertion, so that its C-terminus is cytosolic (N_cyto_-C_cyto_). We show that a mutant form of ZMPSTE24, ZMPSTE24^E336A^, traps and stabilizes this altered topological isoform of IFITM3 and that the N_cyto_-C_cyto_ isoform can also be observed in the absence of ZMPSTE24^E336A^ when the ubiquitin-proteasome system is inhibited. Notably the species of IFITM3 bound to ZMPSTE24^E336A^ is hypo-palmitoylated. We propose that ZMPSTE24 may act as a quality control factor for IFITM3 (Fig. 9). Low palmitoylation may predispose IFITM3 to undergo TM span inversion by ZMPSTE24. Alternatively, ZMPSTE24 may select the aberrant inverted isoform of IFITM3 for re-inversion and if this rescue step fails, then IFITM3 can be degraded by the ubiquitin-proteasome system. The ZMPSTE24^E336A^ “trap” mutant studied here may sequester the inverted IFITM3 isoform, preventing either reinversion and degradation, thereby revealing the presence of this otherwise transient species of IFITM3.

**Figure 9.**
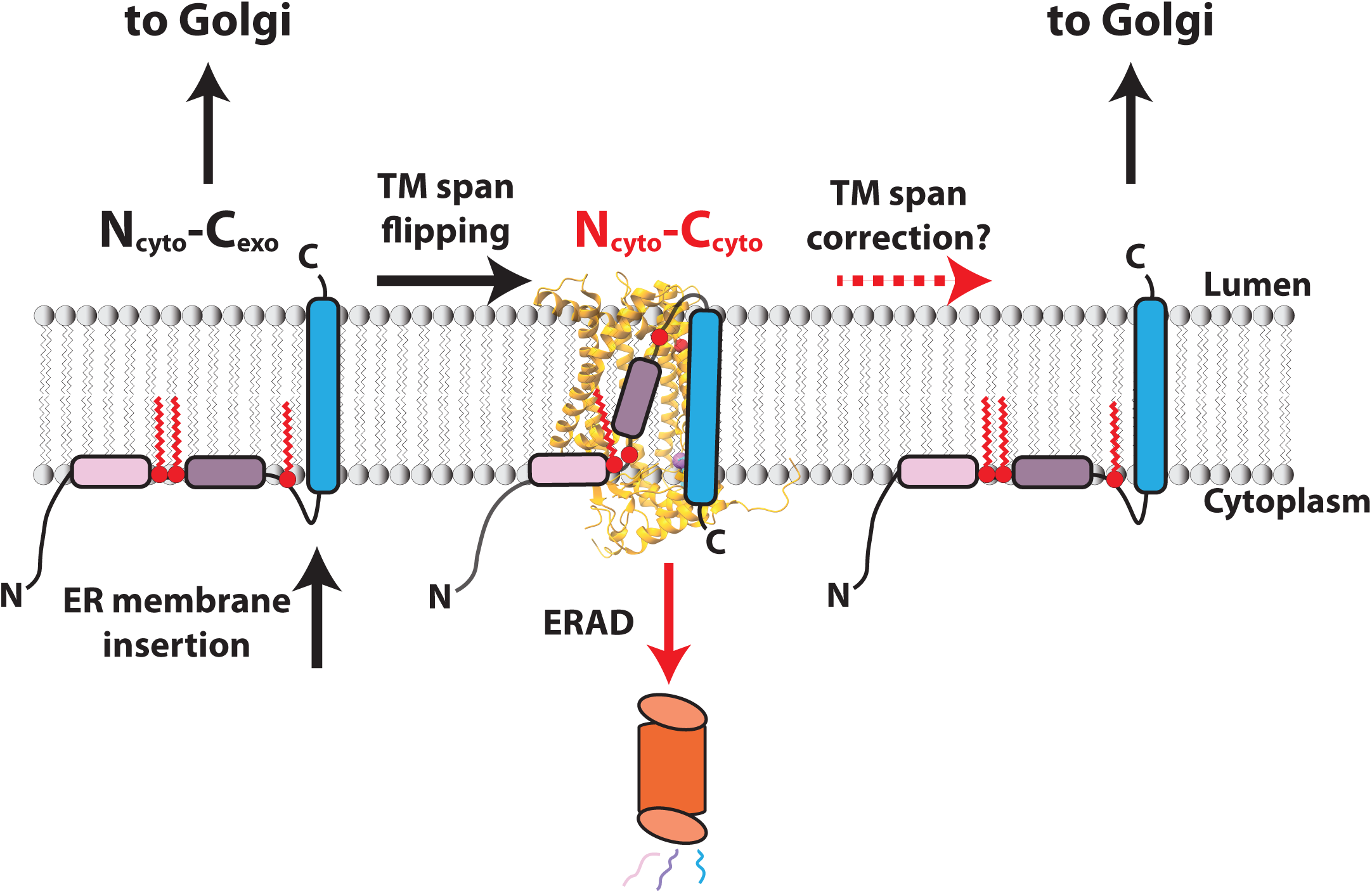
Model for ZMPSTE24’s potential role(s) in the quality control of IFITM3 topology. IFITM3 is initially inserted into the ER membrane in the N_cyto_-C_exo_ topology. Under-palmitoylation of IFITM3 may cause the TM span to flip in the membrane, possibly mediated by ZMPSTE24, resulting in the N_cyto_-C_cyto_ topology. ZMPSTE24 may instead assist in correction of the TM span of IFITM3, allowing trafficking to the Golgi and ultimately the plasma membrane or endosomes where it functions as a viral restriction factor. If not rectified, the N_cyto_-C_cyto_ IFITM3 altered topological form undergoes ubiquitination, membrane extraction and degradation by the proteasome (ERAD). The ZMPSTE24^E336A^ mutant may trap the normally transient inverted form of IFITM3 shown in the middle.

The observation of an inverted TM span for IFITM3 contradicts the long-accepted cell biological dogma that the orientation of a TM span is established during synthesis and remains fixed throughout a protein’s lifetime. However, a growing number of exceptions to this rule have emerged in recent years, including reports of post-translational inversion of TM spans in VAP-B, TM4SF20, and ABCG2 by incompletely understood mechanisms (55–57). In addition, dislocation of mislocalized mitochondrial or ER tail-anchored proteins by the AAA+ proteins with ATPase activity, Msp1/ATAD1 and Spf1/ATP13A1, respectively, maintain organelle identity and allow mislocalized tail-anchored proteins another attempt to properly insert in the correct membrane (58, 59). ATP13A1 is also involved as a determinant in the topology of ABCG2, a multispanning integral membrane protein (55). Since ZMPSTE24 is not an ATPase, nor is it currently known to interact with one, it is unclear how the energy requirement for inverting or reinverting IFITM3’s TM span would be fulfilled. Although speculative, the large volume of the ZMPSTE24 intramembrane chamber is well-suited to accommodate hydrophilic regions adjacent to a TM span that would need to pass from one side of the lipid bilayer to the other (1–3).

A role in protein topology has previously been proposed for the yeast ZMPSTE24 homolog, Ste24 (19, 60). In that study, the authors used an engineered single-spanning TM reporter to select yeast mutants with an altered topology, despite strong topogenic signals encoded within the reporter protein. Mutations in two genes, *STE24* and *SPF1*, were found to increase the pool of the inverse topology of the reporter. Although unknown at the time, it is now well-established, as mentioned above, that the human Spf1 homolog, ATP13A1, functions as a TM span dislocase, ejecting from the ER membrane any mis-localized tail-anchored proteins or mis-inserted TM spans (52, 55, 58, 61). Notably, both ZMPSTE24 and ATP13A1 were identified as high-confidence interactors of IFITM3-Flag (22), suggesting both may function in maintaining the correct topology of IFITM3 during biogenesis. A potential role in yeast for both Ste24/ZMPSTE24 and Spf1/ATP13A1 in maintaining ER protein homeostasis is underscored by the strong activation of the unfolded protein response in *ste24Δ* and *spf1Δ* yeast mutants (20). Likewise, yeast *ste24Δ* mutants show strong growth defects when combined with mutations in ERAD machinery genes (e.g. *ubc7Δ*, *cue1Δ* and *hrd1Δ*) (17, 18), suggesting the ERAD pathway and Ste24 may converge on common substrates, including aberrant topological isoforms of TM proteins.

The use of catalytically inactive enzymes, such as ZMPSTE24^E336A^ that can trap and reveal binding partners, has been a successful strategy for identifying enzyme clientele and proteolytic substrates. For example, mutations that reduce the activities of the ATPases TRC40 and ATP13A1, involved in tail-anchored protein insertion and removal from the ER membrane, respectively, uncovered numerous putative and validated substrates (52, 61, 62). Similarly, active site mutations in the prokaryotic proteases FtsH and ClpXP enabled identification of several physiological substrates that had eluded capture by wild-type enzymes (63–65). However, it is important to note that mere loss of catalytic activity is not sufficient for IFITM3 trapping by ZMPSTE24. Mutations in zinc-coordinating residues (H335A, H339A, and E415A), which abolish prelamin A cleavage, did not enhance binding and, in fact, disrupted the trapping capability of ZMPSTE24^E336A^ (Fig. 2A), possibly because they impact an important structural feature. Substrates with transmembrane spans, such as the IFITMs, may enter the ZMPSTE24 chamber through lateral diffusion within the lipid bilayer, passing between the transmembrane segments of ZMPSTE24. The spacious interior chamber may provide a pathway for adjacent hydrophilic domains that would otherwise be unable to traverse the membrane. The mechanism by which mutation of the catalytic glutamate stabilizes the interaction between ZMPSTE24 and IFITM3 is not currently understood. However, it is possible that it is simply the change of a charged to a nonpolar residue (Glu to Ala) that enhances ZMPSTE24’s binding affinity.

Identifying additional substrates and binding partners of ZMPSTE24 is a critical step in understanding both its physiological roles and the binding determinants for these interactions. A recent large-scale proteome-based screen identified seven proteins that interact with ZMPSTE24, six of which are membrane proteins with a single TM span that localize to the ER or ER-proximal vesicles (66). Notably, four of these proteins (VAP-A, VAP-B, STX8, and ANKRD46) are tail-anchored proteins, while PGRMC1 and CANX are membrane proteins with a single TM span located near the N- and C-termini, respectively. PGRMC1 has been reported to exist in both N_cyto_-C_exo_ and N_exo_-C_cyto_ topologies (67–69), although it remains unclear whether there could be a topological inversion of PGRMC1 during biosynthesis or post-insertion. The ZMPSTE24^E336A^ trapping mutant may prove useful in future studies to validate these interactors and identify new client proteins for ZMPSTE24.

IFITM3 has long been appreciated to play a role in viral defense; when it is overexpressed cells become resistant to many enveloped viruses and when knocked down or deleted, cells are sensitized to viral infection (34, 70, 71). ZMPSTE24, which interacts with the IFITMs, was itself shown more recently to have similar properties, namely its overexpression promoted viral defense (and notably catalytic activity is dispensable for this role) and knocking down ZMPSTE24 in cells or infecting *Zmpste24^-/-^*mice sensitized them to viral infection (22, 24, 25). IFITMs are thought to promote membrane stiffness, thereby preventing viral-host cell membrane fusion (38, 45). Exactly how ZMPSTE24 defends against viral infection remains an important unanswered question. It could act through its role shown here in influencing IFITM3 topology and/or quality control. Alternatively, ZMPSTE24 may have a more generalizable role in membrane integrity or properties by affecting membrane proteins influencing membrane fluidity. Further studies will be important to distinguish these possibilities. A deeper understanding of ZMPSTE24’s mechanistic role in terms of IFITM3 and viral defense could ultimately lead to the development of novel anti-viral therapeutic options.

### Experimental Procedures

#### Cell lines, Plasmids and Antibodies

Knockout of *ZMPSTE24* in HEK293T was done using CRISPR-Cas9 lentiviral transduction followed by limited dilution and confirmed by Western blotting. The topology reporter cell line (Flp-In 293 T-Rex cells containing mNeon Green(1–10)_cyto_ and mCherry(1–10)_ER_) was provided by Susan Shao (Harvard Medical School). Stable integration of IFITM3 constructs into the Flp-In locus was done by co-transfecting pcDNA5 FRT/TO plasmids with pOG44 (1:3 ratio) using Lipofectamine 3000 according to the manufacturer’s instructions and selected with 50 μg/ml Hygromcin B, 10 μg/ml blasticidin and 1 μg/ml puromycin.

Plasmids used in this study are listed in Table 1. Plasmids designated in the text and figures as Flag-tagged contain 2 copies of Flag (2XFlag). 2XFlag-ZMPSTE24 (pSM3914), 2XFlag-ZMPSTE24^E336A^ (pSM3915) and myc-tagged IFITM plasmids were reported previously (24). 2XFlag-IFITM3 was constructed by replacing the *ZMPSTE24* ORF from pSM3914 with a PCR fragment containing the *IFITM3* gene by HiFi Assembly (New England Biolabs). Mutations in *ZMPSTE24* and *IFITM3* plasmids were introduced by PCR using mutagenic oligonucleotides and recombinational cloning using NEB HiFi Assembly (New England Biolabs). pSM4117 was constructed by replacing the *Flag-ATP13A1* gene from pcDNA5 FRT/TO Flag-ATP13A1 (61) with a PCR fragment containing *3Xmyc-LMNA*(*431-664*). Opsin-tagged IFITM3 plasmids pSM3917 and pSM4004 were constructed by placing nucleotides corresponding to an opsin glycosylation tag (GPNFYVPFS**N**KTG) before the stop codon (pSM3917) or after codon 40 (pSM4004) of the *IFITM3* gene, using pSM3867 as template. Topology test plasmids pSM4085 (β11-IFITM3) and pSM4121 (IFITM3-β11) were constructed using pcDNA5 FRT/TO BFP-P2A-β11-Sec61β (52) as template, replacing the *SEC61β* gene with IFITM3 tagged at either the N-terminus or C-terminus.

**Table 1.** Plasmids used in this study.

| **Plasmid** | **Protein** | **Source** |
| --- | --- | --- |
| pSM3867 | myc-IFITM3 | Addgene #58461 |
| pSM3873 | myc-IFITM1 | This study |
| pSM3874 | myc-IFITM2 | This study |
| pSM3875 | HA-IFITM3 | This study |
| pSM3910 | 10His-3HA-ZMPSTE24-E336A | This study |
| pSM3913 | 2XFlag-IFITM3 | This study |
| pSM3914 | 2XFlag-ZMPSTE24 | (24) |
| pSM3915 | 2XFlag-ZMPSTE24-E336A | (24) |
| pSM3917 | myc-IFITM3-opsin-C | This study |
| pSM3920 | myc-IFITM3-C105A | This study |
| pSM3969 | 2XFlag-ZMPSTE24-C324A/E336A | This study |
| pSM3977 | 2XFlag-ZMPSTE24-E336A-cysteineless | This study |
| pSM3978 | pCMV-empty | This study |
| pSM3981 | myc-IFITM3-C71A | This study |
| pSM3982 | myc-IFITM3-C72A | This study |
| pSM3986 | 2XFlag-ZMPSTE24-C109A, E336A | This study |
| pSM3987 | 2XFlag-ZMPSTE24-C176A, E336A | This study |
| pSM3988 | 2XFlag-ZMPSTE24-C359A, E336A | This study |
| pSM3989 | 2XFlag-ZMPSTE24-C406A, E336A | This study |
| pSM3994 | 3Xmyc-ZMPSTE24 | This study |
| pSM3995 | 3Xmyc-ZMPSTE24-E336A | This study |
| pSM4004 | myc-IFITM3-opsin40 | This study |
| pSM4026 | myc-IFITM3-C72A, C105A | This study |
| pSM4059 | 2XFlag-ZMPSTE24-E336A, C359 only | This study |
| pSM4061 | pOG44 FLP | Thermofisher Scientific |
| pSM4066 | ASGR1-mCherry11-GFP11-P2A-BFP | (52) |
| pSM4069 | 2XFlag-ZMPSTE24-H339A | This study |
| pSM4070 | 2XFlag-ZMPSTE24-E415A | This study |
| pSM4080 | 2XFlag-ZMPSTE24-E336A, E415A | This study |
| pSM4085 | BFP-P2A-mCherry11-GFP11-IFITM3 | This study |
| pSM4095 | 2XFlag-ZMPSTE24-H335A | This study |
| pSM4117 | 3Xmyc-LMNA(431-664) | This study |
| pSM4120 | 2XFlag-ZMPSTE24-E336A, H339A | This study |
| pSM4121 | BFP-P2A-IFITM3-mCherry11-GFP11 | This study |
| pSM4122 | BFP-P2A-IFITM3-C71A-mCherry11-GFP11 | This study |

Anti-Flag (Sigma, F1804) and anti-DYKDDDK (FG4R) DyLight^TM^ 680 (Thermofisher, MA1-91878-D680) were used for Western blotting at 1:5000 and 1:1000, respectively. Anti-myc (71D10) rabbit mAb (Cell Signaling Technology, 2278S) was used at 1:2000 dilution. Anti-IFITM1 (Proteintech, 11727-3-AP) was used at 1:1000. Anti-IFITM2 (Proteintech, 12769-1-AP) was used at 1:2000. Anti-IFITM3 (Proteintech, 11714-1-AP) was used at 1:10,000. Goat anti-mouse (Licor, 925-68070), goat anti-rat (Licor, 925-68076) and goat anti-rabbit (Licor, 925-32211) secondary antibodies were used at 1:15,000 dilution.

#### Co-IP, EndoH/PNGase F treatment and Western blotting

Co-IP experiments were done as previously described (24). Briefly, HEK293T cells were seeded in 6-well tissue culture plates and transfected with 300 ng plasmid (typically 200 ng Flag-ZMPSTE24 with 100 ng myc-IFITM3 plasmids) using Effectene transfection reagent (Qiagen) according to the manufacturer’s instructions. After 48 hrs, cells were collected, washed with cold PBS and resuspended in cold lysis buffer (50 mM HEPES pH 7.5, 150 mM NaCl, 1% Triton X-100, 1X protease inhibitor cocktail (Roche) and 0.5 mM PMSF. Cells were solubilized for 45 min at 4°C by end-over-end rotation. Lysates were centrifuged at 22k x g for 15 minutes and again at 100k x g for 30-60 minutes, and then the soluble fraction was mixed end-over-end with 15 μl of anti-Flag magnetic agarose (Sigma) for 2 hrs. The beads were washed twice with cold lysis buffer and once with cold PBS. Proteins were eluted using 1X SDS sample buffer (Biorad), heated at 65°C for 15 minutes and resolved by SDS-PAGE (4-20% Criterion TGX; Biorad). β-mercaptoethanol was added to the sample buffer to a final concentration of 5% prior to heating for reducing SDS-PAGE, but omitted for non-reducing SDS-PAGE. For de-glycosylation experiments, immunoprecipitated proteins were eluted from anti-Flag beads using 10mM sodium phosphate pH 7.5, 0.5% SDS before treatment with EndoH (NEB) or PNGase F (NEB) according to the manufacturer’s instructions. Proteins were transferred to nitrocellulose using the TransBlot Turbo (Biorad) Western transfer apparatus and blocked with PBS containing 0.1% Tween-20 (PBS-T) with 5% nonfat dry milk for 30-60 minutes at room temperature. Blots were incubated with the indicated antibodies in PBS-T overnight at 4°C, washed three times with PBS-T, and incubated with the appropriate secondary antibodies. After three PBS-T washes, the blot was scanned on the Odyssey CLx (Licor) and analyzed using Image Studio Lite software (Licor). For experiments comparing detergent-dependent disulfide bond formation (Fig. S2), digitonin (Millipore), n-Dodecyl β-D-maltoside (DDM, Millipore) and Triton X-100 (Biorad) were used at 1% final concentration in lysis buffer.

#### Metabolic labeling and click chemistry for palmitoylation

To examine palmitoylation of ZMPSTE24^E336A^-trapped IFITM3, bio-orthogonal labeling using 17-octadecynoic acid (17-ODYA; Cayman Chemical) and click chemistry was done essentially as described (50), with minor modifications. HEK293T cells were co-transfected with HA-ZMPSTE24^E336A^/Flag-IFITM3 or HA-IFITM3/Flag-IFITM3. The next day, the medium was removed and replaced with DMEM containing 5% charcoal-stripped FBS (Thermofisher) for 2 hrs. Labeling was initiated by adding 17-ODYA (30 μM) in DMEM with 5% charcoal-stripped FBS for 3 hours. Cells were collected, washed with PBS, and lysed in 50mM triethanolamine (TEA) pH 7.5, 150mM NaCl, 1% Triton X-100 containing 1X EDTA-free protease inhibitor cocktail (Roche) for 30 min at 4°C. The lysate was centrifuged at 22k x g for 15 min at 4°C, and the supernatant was mixed end-over-end with 15 μl anti-HA magnetic agarose (Sigma) for 2 hours at 4°C. The beads were washed twice in lysis buffer, and once with cold PBS. Immune complexes were released by adding 50mM TEA pH 7.5, 150mM NaCl, 1% SDS, a portion of which was resolved by SDS-PAGE and Western blotting to confirm the co-IP. The rest of the eluate was diluted in lysis buffer so that the SDS was 0.05% and then immunoprecipitated with anti-Flag magnetic agarose for 2 hours. The beads were washed three times in modified RIPA buffer (50mM TEA pH 7.5, 150mM NaCl, 1% Triton X-100, 1% sodium deoxycholate, 0.1% SDS). On-bead click chemistry was initiated by adding 50 μl of click chemistry reaction mix (1mM TCEP, 1mM CuSO_4_-5H_2_O, 100 uM TBTA, 100 uM IRDye800 CW azide (Licor) in PBS) for 1 hour at RT. The beads were washed 3 times with RIPA buffer, and proteins were eluted with 25 ul 50mM TEA pH 7.5, 150mM NaCl, 4% SDS heated at 37°C for 10 min. The eluate was added to an equal volume of Biorad 4X sample buffer containing 10% β-ME (5% final) and heated at 65°C for 10 minutes. Samples were resolved by 4-20% SDS-PAGE (Biorad) and transferred to nitrocellulose. The nitrocellulose was blocked with PBS-T containing 5% nonfat dry milk, probed with mouse anti-Flag (1:10,000 dilution), followed by goat anti-mouse IRDye680 secondary antibody. The blot was imaged on a Licor Oddysey CLx, which detected both the Flag proteins (red) and azide-modified IRDye800 proteins (green).

#### Fluorescence microscopy and flow cytometry

For fluorescence microscopy, topology reporter cell lines propagated in 24-well or 6-well tissue culture plates were transfected with empty vector (pSM3978), 3Xmyc-ZMPSTE24 (pSM3994) or 3Xmyc-ZMPSTE24^E336A^ (pSM3995) using Effectene transfection reagent. The following day, doxycycline (200 ng/ml) was added to induce topology reporter proteins for 24 hours. Images were collected on a Molecular Devices ImageXpress Micro XLS Widefield high-Content Analysis System.

For fluorescence flow cytometry, cells were grown in 12-well or 24-well tissue culture plates, transfected and induced as described above. Cells were trypsinized, collected by centrifugation (800 x g) and washed with cold PBS. Cells were filtered through a 35 μM mesh filter prior to data collection on an Attune NxT flow cytometer. Data were analyzed using the free online software Floreada (floreada.io). Only single cells expressing high BFP fluorescence were used for data analysis. Average GFP:BFP ratios were used to determine fold changes between cell lines or treatments. For drug treatments, cells were induced with doxycycline for 24 hours prior to addition of the proteasome inhibitor bortezomb (1μM), the E1 ubiquitin activating enzyme inhibitor MLN4273 (1μM; Cayman Chemical 1450833-55-2), the p97/VCP inhibitor NMS-873 (5 μM; Ape Bio B2168) or DMSO control for 4 hours, followed by flow cytometry.

#### Statistical analysis

Statistical analyses were performed using GraphPad Prism. Group mean comparisons were conducted using one-way ANOVA, followed by Dunnett’s multiple comparisons test to identify specific group differences.

## Data availability

All data are included in the main text and Supporting Information.

## Supporting Information

This article containing Supporting Information.

## Acknowledgments

We thank Michael McKenna and Susan Shao (Harvard Medical School) for sharing the topology reporter cell lines and plasmids. We thank Elana Shaw and Rohan Panaparambil for help with flow cytometry data collection and data analysis. We thank LaToya Roker and the Johns Hopkins School of Medicine Microscope Facility for assistance with fluorescence microscopy.

Funding and Additional Information. This study was funded by an NIH grant R35GM127073 from NIGMS to S.M.

## Figure Legends

**Figure S1.**
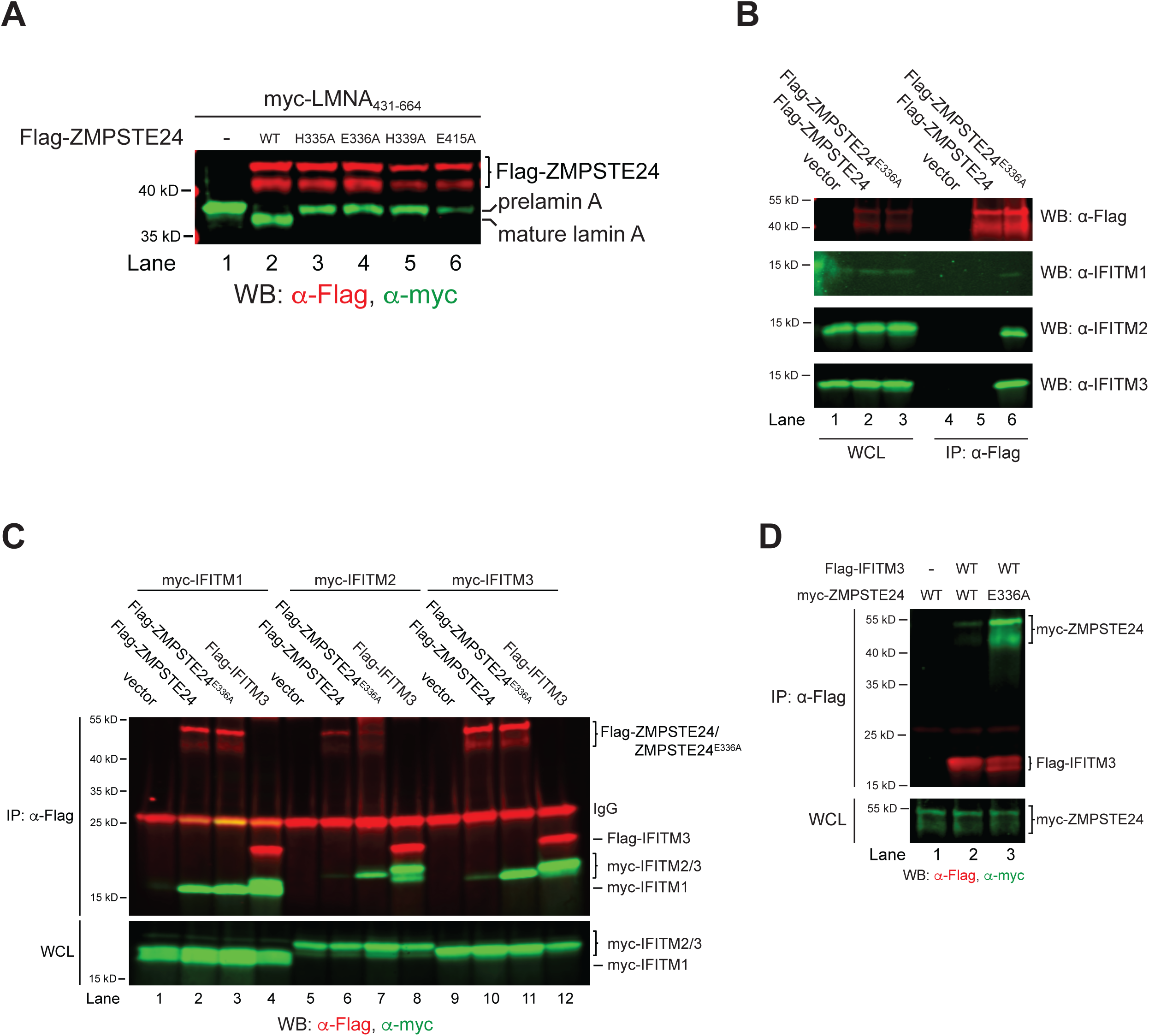
ZMPSTE24^E336A^ shows enhanced interactions with IFITM1, 2, and 3 proteins. (A) Assessing loss of catalytic activity for ZMPSTE24 HExxH…E mutants. Flag-tagged wild-type and mutant ZMPSTE24 plasmids were co-transfected with a myc-tagged prelamin A plasmid (myc-LMNA_(431-664)_) in *ZMPSTE24* knockout HEK293 cells to evaluate ZMPSTE24 proteolytic activity. Flag-ZMPSTE24, prelamin A and its cleaved product, mature lamin A, were detected by Western blotting and are indicated. All active site mutants block cleavage of the prelamin A construct. (B) Co-IP’s of endogenous IFITMs by ZMPSTE24 and ZMPSTE24^E336A^. HEK293 cells transfected with vector, Flag-ZMPSTE24 or Flag-ZMPSTE24^E336A^ were induced with 150 units of interferon β for 18 hours prior to immunoprecipitation with anti-Flag agarose. Whole cell lysate (WCL) and IP’d proteins were resolved by SDS-PAGE analyzed by Western blotting with anti-Flag (ZMPSTE24), anti-IFITM1, IFITM2 or IFITM3 antibodies. (C) Comparing co-IPs of IFITMs by Flag-ZMPSTE24^E336A^ and Flag-IFITM3. Cells transfected with vector, Flag-ZMPSTE24, Flag-ZMPSTE24^E336A^ or Flag-IFITM3 and the indicated myc-tagged IFITM proteins were subjected to co-IP as described. Positions of ZMPSTE24 and IFITM proteins are indicated. Lanes 9-12 were used to make Figure 2C. (D) Flag-tagged IFITM3 co-IPs myc-ZMPSTE24^E336A^ more efficiently than wild-type myc-ZMPSTE24. HEK293 cells were transfected with Flag-IFITM3 and myc-ZMPSTE24 or myc-ZMPSTE24^E336A^. Proteins were co-IP’d with anti-Flag agarose prior to SDS-PAGE and Western blotting. Positions of Flag-IFITM3 and myc-ZMPSTE24 proteins are indicated.

**Figure S2.**
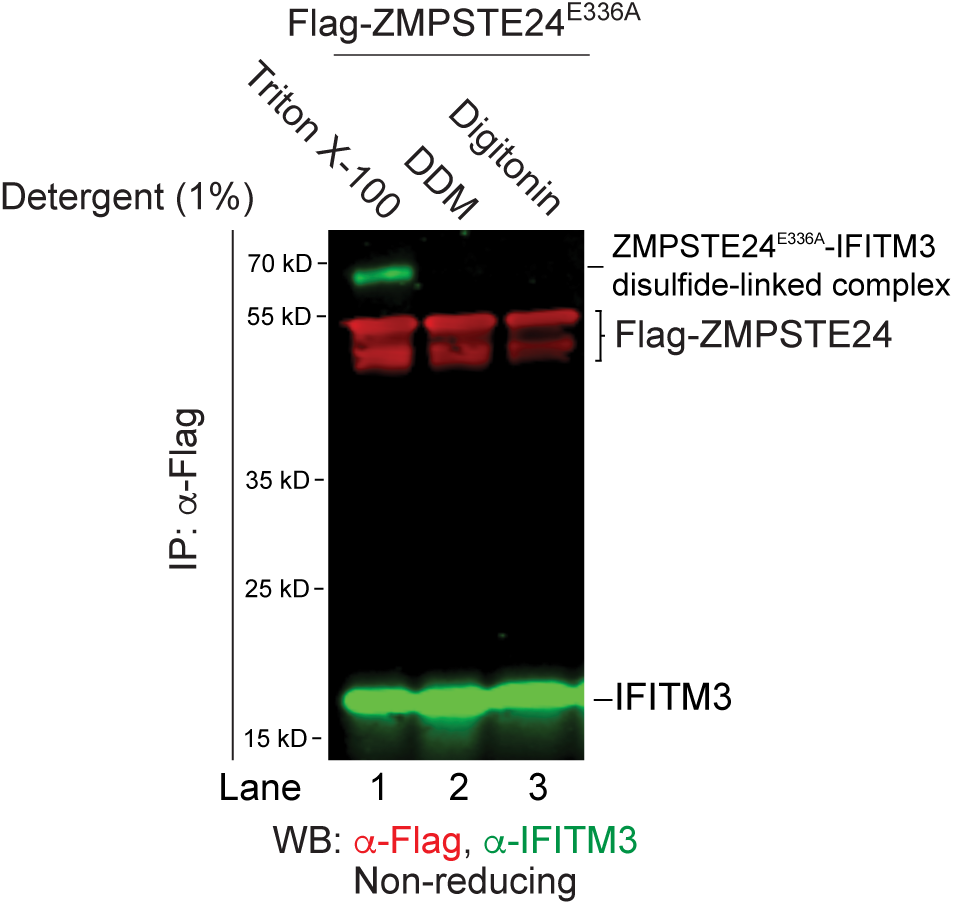
Detergent-dependence of the ZMPSTE24^E336A^-IFITM3 intermolecular disulfide complex. Cells expressing Flag-ZMPSTE24^E336A^ were induced with interferon β for 18 hrs. Cells were lysed in buffer containing 1% Triton X-100, 1% dodecyl maltoside (DDM) or 1% digitonin prior to immunoprecipitation with anti-Flag agarose beads. Proteins were resolved by non-reducing SDS-PAGE and analyzed by Western blotting with anti-Flag and anti-IFITM3 antibodies. Positions of IFITM3, Flag-ZMPSTE24^E336A^ and the intermolecular disulfide-linked complex are shown.

**Figure S3.**
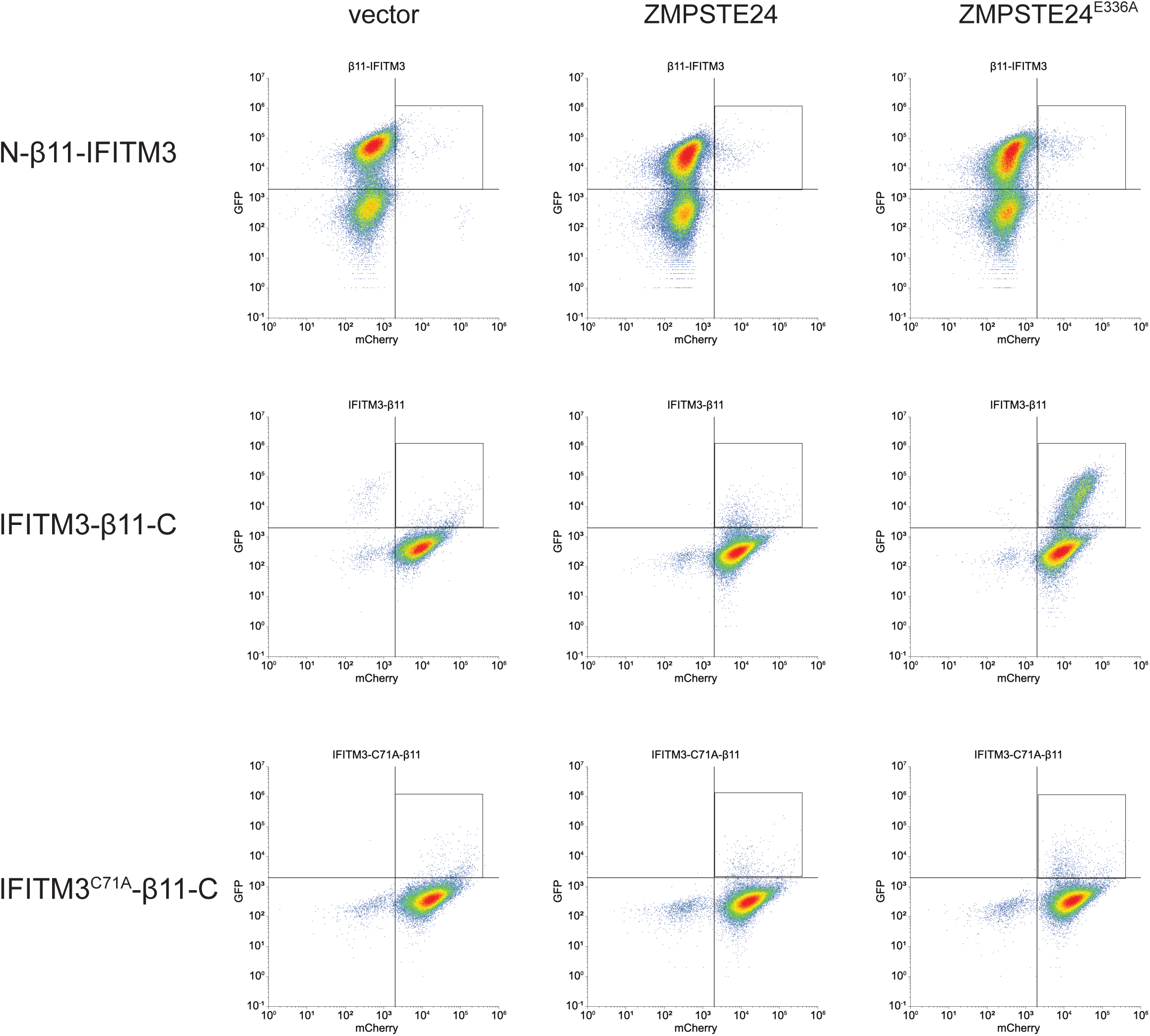
Flow cytometry shows IFITM3 is normally N_cyto_-C_exo_, but expression of ZMPSTE24^E336A^ yields a subpopulation with a cytosolic C-terminus (N_cyto_-C_cyto_). 293 TREx topology reporter cells stably transfected with N-β11-IFITM3, IFITM3-β11-C or IFITM3^C71A^-β11 were transfected with plasmids empty vector, ZMPSTE24 or ZMPSTE24^E336A^ and analyzed by flow cytometry. A box in the right upper quadrant is shown to highlight cells with high GFP and mCherry fluorescent signals. IFITM3-C71A-β11, which does not interact with IFITM3, does not exhibit a subpopulation with a cytosolic C-terminus in the presence of ZMPSTE24^E336A^. Data shown here is representative of the six trials used for quantification shown in Fig. 6D.

**Figure S4.**
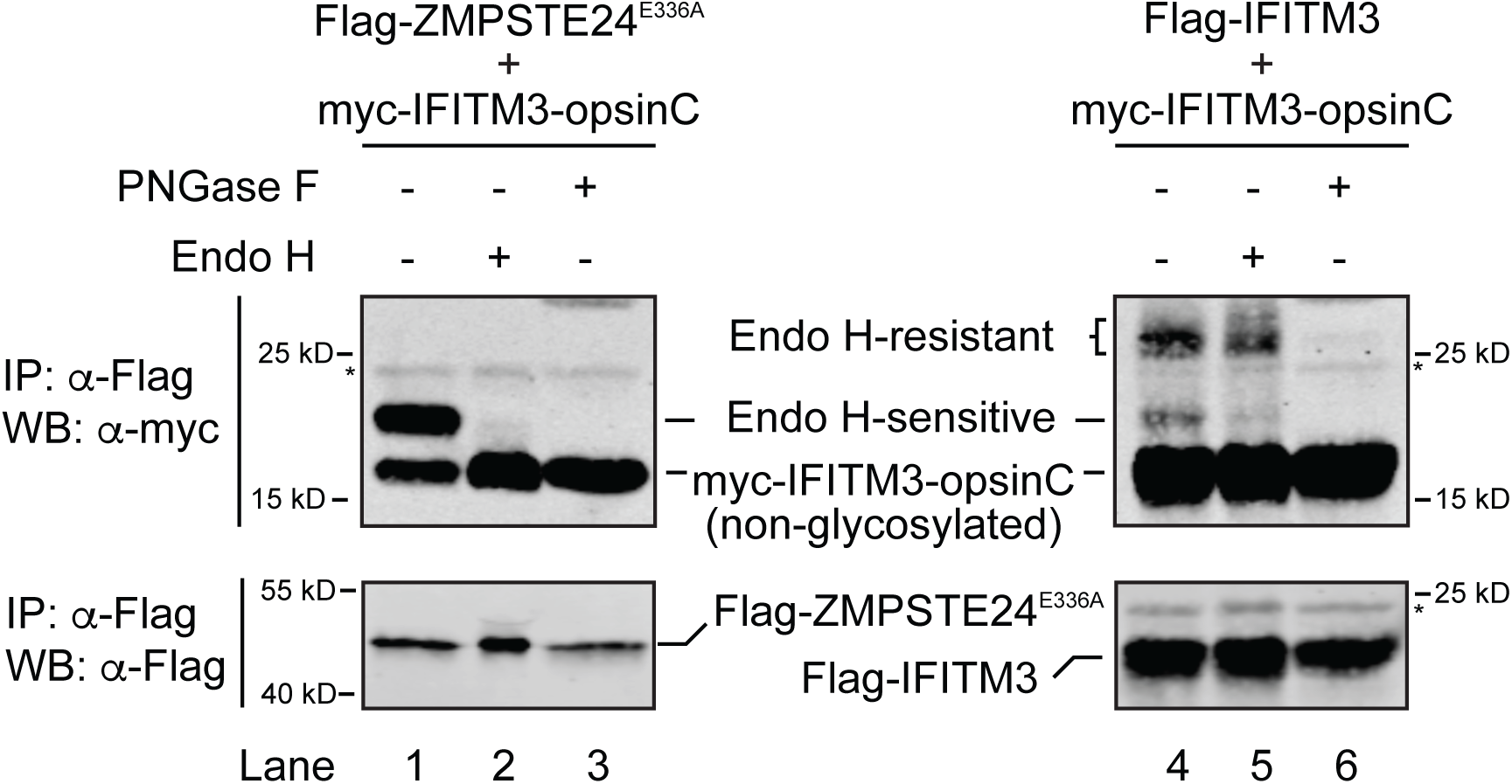
IFITM3-opsinC that co-purifies with ZMPSTE24^E336A^ is EndoH-sensitive, while IFITM3-opsinC recovered in IFITM3 homo-oligomers is EndoH-resistant, consistent with Golgi glycosylation of the oligomer-associated species but not the ZMPSTE24^E336A^-bound species. HEK293 cells were transfected with Flag-tagged ZMPSTE24^E336A^ and myc-IFITM3-opsinC (left panels) or Flag-tagged IFITM3 and myc-IFITM3-opsinC (right panels). Proteins were immunoprecipitated with anti-Flag agarose and then mock treated or treated with EndoH or PNGase prior to SDS-PAGE and western blotting. Positions of the glycosylated isoforms of IFITM3-opsinC are indicated. IFITM3-opsinC co-precipitated by ZMPSTE24^E336A^ was EndoH-and PNGase-sensitive. A glycosylated species of IFITM3-opsinC co-precipitated by Flag-IFITM3 was EndoH-resistant (compare lanes 4 and 5), indicating it had been modified by Golgi-localized glycosyltransferases.

**Figure S5.**
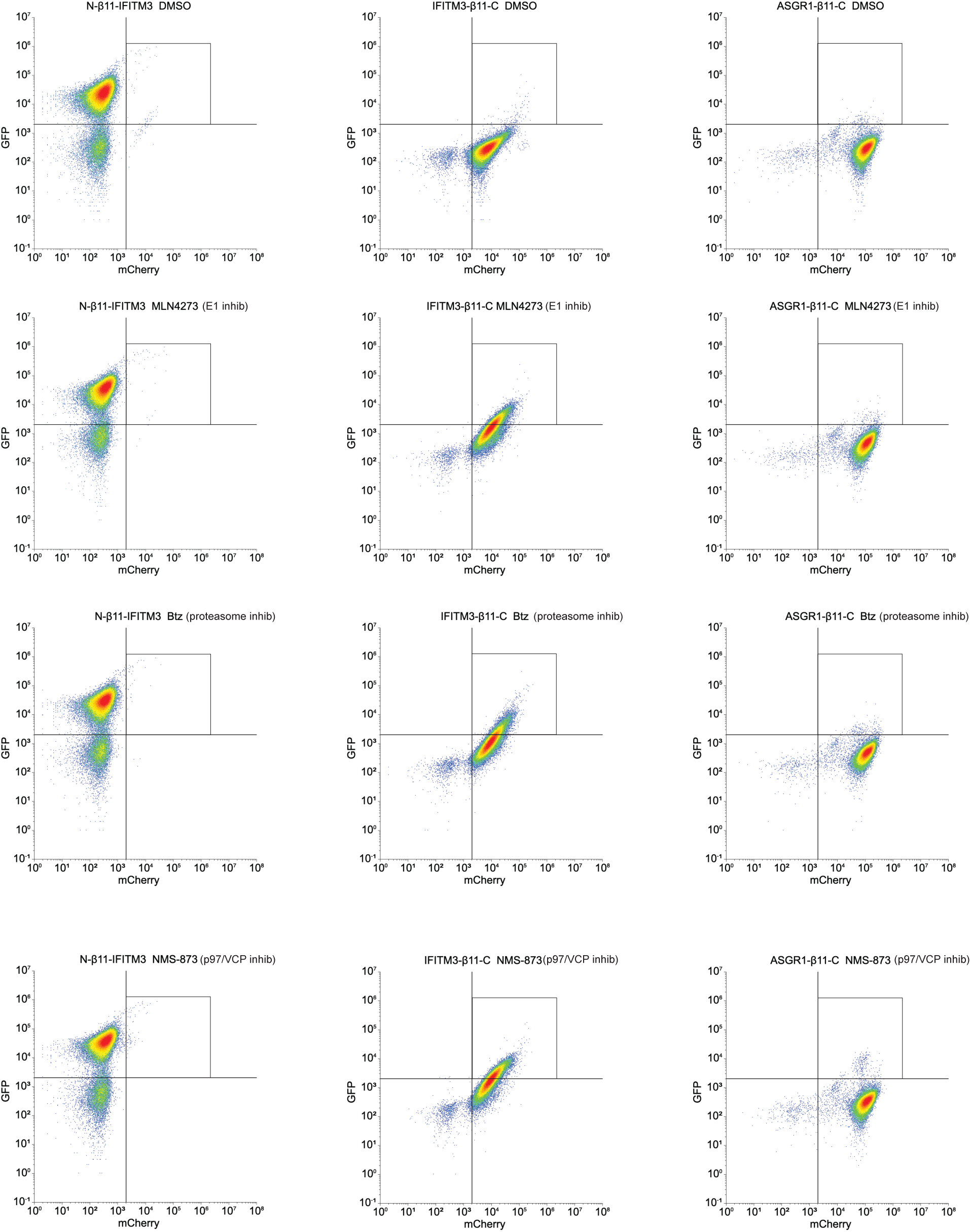
ERAD inhibitors stabilize the IFITM3 C_cyto_ topological isoform. Flow cytometry data from HEK293 cells expressing the indicated IFITM3 constructs were treated with vehicle (DMSO), a ubiquitin E1 inhibitor (E1 inhib), the proteasome inhibitor Bortezomib (Btz), or the p97/VCP inhibitor NMS-873 for 4 hrs. A box in the upper right quadrant highlights cells with elevated GFP and mCherry signals.

## Notes

### Competing Interest Statement

The authors have declared no competing interest.

## References

1. Clark, K. M., Jenkins, J. L., Fedoriw, N., and Dumont, M. E. (2017) Human CaaX protease ZMPSTE24 expressed in yeast: Structure and inhibition by HIV protease inhibitors Protein Sci 26, 242–257 10.1002/pro.3074

2. Pryor, E. E., Jr., Horanyi, P. S., Clark, K. M., Fedoriw, N., Connelly, S. M., Koszelak-Rosenblum, M. et al. (2013) Structure of the integral membrane protein CAAX protease Ste24p Science 339, 1600–1604 339/6127/1600 [pii] 10.1126/science.1232048

3. Quigley, A., Dong, Y. Y., Pike, A. C., Dong, L., Shrestha, L., Berridge, G. et al. (2013) The structural basis of ZMPSTE24-dependent laminopathies Science 339, 1604–1607 10.1126/science.1231513

4. Spear, E. D., and Michaelis, S. (2025) Mammalian zinc metalloprotease ZMPSTE24 and yeast Ste24 In Handbook of Proteolytic Enzymes: Metallopeptidases, 537–549

5. Bergo, M. O., Gavino, B., Ross, J., Schmidt, W. K., Hong, C., Kendall, L. V. et al. (2002) Zmpste24 deficiency in mice causes spontaneous bone fractures, muscle weakness, and a prelamin A processing defect Proc Natl Acad Sci U S A 99, 13049–13054 10.1073/pnas.192460799

6. Dittmer, T. A., and Misteli, T. (2011) The lamin protein family Genome Biol 12, 222 gb-2011-12-5-222 [pii] 10.1186/gb-2011-12-5-222

7. Pendas, A. M., Zhou, Z., Cadinanos, J., Freije, J. M., Wang, J., Hultenby, K. et al. (2002) Defective prelamin A processing and muscular and adipocyte alterations in Zmpste24 metalloproteinase-deficient mice Nat Genet 31, 94–99 10.1038/ng871

8. Boyartchuk, V. L., Ashby, M. N., and Rine, J. (1997) Modulation of Ras and a-factor function by carboxyl-terminal proteolysis Science 275, 1796–1800 10.1126/science.275.5307.1796

9. Boyartchuk, V. L., and Rine, J. (1998) Roles of prenyl protein proteases in maturation of Saccharomyces cerevisiae a-factor Genetics 150, 95–101, http://www.ncbi.nlm.nih.gov/pubmed/9725832

10. Fujimura-Kamada, K., Nouvet, F. J., and Michaelis, S. (1997) A novel membrane-associated metalloprotease, Ste24p, is required for the first step of NH2-terminal processing of the yeast a-factor precursor J Cell Biol 136, 271–285, http://www.ncbi.nlm.nih.gov/entrez/query.fcgi?cmd=Retrieve&db=PubMed&dopt=Citation&list_uids=9015299

11. Schmidt, W. K., Tam, A., and Michaelis, S. (2000) Reconstitution of the Ste24p-dependent N-terminal proteolytic step in yeast a-factor biogenesis J Biol Chem 275, 6227–6233, https://www.ncbi.nlm.nih.gov/pubmed/10692417

12. Tam, A., Nouvet, F. J., Fujimura-Kamada, K., Slunt, H., Sisodia, S. S., and Michaelis, S. (1998) Dual roles for Ste24p in yeast a-factor maturation: NH2-terminal proteolysis and COOH-terminal CAAX processing J Cell Biol 142, 635–649, https://www.ncbi.nlm.nih.gov/pubmed/9700155

13. Tam, A., Schmidt, W. K., and Michaelis, S. (2001) The multispanning membrane protein Ste24p catalyzes CAAX proteolysis and NH2-terminal processing of the yeast a-factor precursor J Biol Chem 276, 46798–46806 10.1074/jbc.M106150200

14. Barrowman, J., Wiley, P. A., Hudon-Miller, S. E., Hrycyna, C. A., and Michaelis, S. (2012) Human ZMPSTE24 disease mutations: residual proteolytic activity correlates with disease severity Hum Mol Genet 21, 4084–4093 10.1093/hmg/dds233

15. Spear, E. D., Alford, R. F., Babatz, T. D., Wood, K. M., Mossberg, O. W., Odinammadu, K. et al. (2019) A humanized yeast system to analyze cleavage of prelamin A by ZMPSTE24 Methods 157, 47–55 10.1016/j.ymeth.2019.01.001

16. Spear, E. D., Hsu, E. T., Nie, L., Carpenter, E. P., Hrycyna, C. A., and Michaelis, S. (2018) ZMPSTE24 missense mutations that cause progeroid diseases decrease prelamin A cleavage activity and/or protein stability Dis Model Mech 11, 10.1242/dmm.033670

17. Costanzo, M., Baryshnikova, A., Bellay, J., Kim, Y., Spear, E. D., Sevier, C. S. et al. (2010) The genetic landscape of a cell Science 327, 425–431 10.1126/science.1180823

18. Usaj, M., Tan, Y., Wang, W., VanderSluis, B., Zou, A., Myers, C. L. et al. (2017) TheCellMap.org: A Web-Accessible Database for Visualizing and Mining the Global Yeast Genetic Interaction Network G3 (Bethesda) 7, 1539–1549 10.1534/g3.117.040220

19. Tipper, D. J., and Harley, C. A. (2002) Yeast genes controlling responses to topogenic signals in a model transmembrane protein Mol Biol Cell 13, 1158–1174 10.1091/mbc.01-10-0488

20. Jonikas, M. C., Collins, S. R., Denic, V., Oh, E., Quan, E. M., Schmid, V. et al. (2009) Comprehensive characterization of genes required for protein folding in the endoplasmic reticulum Science 323, 1693–1697 10.1126/science.1167983

21. Ast, T., Michaelis, S., and Schuldiner, M. (2016) The Protease Ste24 Clears Clogged Translocons Cell 164, 103–114 10.1016/j.cell.2015.11.053

22. Fu, B., Wang, L., Li, S., and Dorf, M. E. (2017) ZMPSTE24 defends against influenza and other pathogenic viruses J Exp Med 214, 919–929 10.1084/jem.20161270

23. Li, S., Fu, B., Wang, L., and Dorf, M. E. (2017) ZMPSTE24 Is Downstream Effector of Interferon-Induced Transmembrane Antiviral Activity DNA Cell Biol 36, 513–517 10.1089/dna.2017.3791

24. Shilagardi, K., Spear, E. D., Abraham, R., Griffin, D. E., and Michaelis, S. (2022) The Integral Membrane Protein ZMPSTE24 Protects Cells from SARS-CoV-2 Spike-Mediated Pseudovirus Infection and Syncytia Formation mBio 13, e0254322 10.1128/mbio.02543-22

25. Stott-Marshall, R. J., and Foster, T. L. (2022) Inhibition of Arenavirus Entry and Replication by the Cell-Intrinsic Restriction Factor ZMPSTE24 Is Enhanced by IFITM Antiviral Activity Front Microbiol 13, 840885 10.3389/fmicb.2022.840885

26. Brass, A. L., Huang, I. C., Benita, Y., John, S. P., Krishnan, M. N., Feeley, E. M. et al. (2009) The IFITM proteins mediate cellular resistance to influenza A H1N1 virus, West Nile virus, and dengue virus Cell 139, 1243–1254 10.1016/j.cell.2009.12.017

27. Chesarino, N. M., Compton, A. A., McMichael, T. M., Kenney, A. D., Zhang, L., Soewarna, V. et al. (2017) IFITM3 requires an amphipathic helix for antiviral activity EMBO Rep 18, 1740–1751 10.15252/embr.201744100

28. John, S. P., Chin, C. R., Perreira, J. M., Feeley, E. M., Aker, A. M., Savidis, G. et al. (2013) The CD225 domain of IFITM3 is required for both IFITM protein association and inhibition of influenza A virus and dengue virus replication J Virol 87, 7837–7852 10.1128/JVI.00481-13

29. McMichael, T. M., Zhang, Y., Kenney, A. D., Zhang, L., Zani, A., Lu, M. et al. (2018) IFITM3 Restricts Human Metapneumovirus Infection J Infect Dis 218, 1582–1591 10.1093/infdis/jiy361

30. Shi, G., Kenney, A. D., Kudryashova, E., Zani, A., Zhang, L., Lai, K. K., et al. (2021) Opposing activities of IFITM proteins in SARS-CoV-2 infection EMBO J 40, e106501 10.15252/embj.2020106501

31. Zhao, X., Sehgal, M., Hou, Z., Cheng, J., Shu, S., Wu, S., et al. (2018) Identification of Residues Controlling Restriction versus Enhancing Activities of IFITM Proteins on Entry of Human Coronaviruses J Virol 92, 10.1128/JVI.01535-17

32. Prelli Bozzo, C., Nchioua, R., Volcic, M., Koepke, L., Kruger, J., Schutz, D., et al. (2021) IFITM proteins promote SARS-CoV-2 infection and are targets for virus inhibition in vitro Nat Commun 12, 4584 10.1038/s41467-021-24817-y

33. Xie, Q., Bozzo, C. P., Eiben, L., Noettger, S., Kmiec, D., Nchioua, R., et al. (2023) Endogenous IFITMs boost SARS-coronavirus 1 and 2 replication whereas overexpression inhibits infection by relocalizing ACE2 iScience 26, 106395 10.1016/j.isci.2023.106395

34. Shi, G., Schwartz, O., and Compton, A. A. (2017) More than meets the I: the diverse antiviral and cellular functions of interferon-induced transmembrane proteins Retrovirology 14, 53 10.1186/s12977-017-0377-y

35. Bailey, C. C., Kondur, H. R., Huang, I. C., and Farzan, M. (2013) Interferon-induced transmembrane protein 3 is a type II transmembrane protein J Biol Chem 288, 32184–32193 10.1074/jbc.M113.514356

36. Weston, S., Czieso, S., White, I. J., Smith, S. E., Kellam, P., and Marsh, M. (2014) A membrane topology model for human interferon inducible transmembrane protein 1 PLoS One 9, e104341 10.1371/journal.pone.0104341

37. Desai, T. M., Marin, M., Chin, C. R., Savidis, G., Brass, A. L., and Melikyan, G. B. (2014) IFITM3 restricts influenza A virus entry by blocking the formation of fusion pores following virus-endosome hemifusion PLoS Pathog 10, e1004048 10.1371/journal.ppat.1004048

38. Guo, X., Steinkuhler, J., Marin, M., Li, X., Lu, W., Dimova, R. et al. (2021) Interferon-Induced Transmembrane Protein 3 Blocks Fusion of Diverse Enveloped Viruses by Altering Mechanical Properties of Cell Membranes ACS Nano 15, 8155–8170 10.1021/acsnano.0c10567

39. Li, K., Markosyan, R. M., Zheng, Y. M., Golfetto, O., Bungart, B., Li, M., et al. (2013) IFITM proteins restrict viral membrane hemifusion PLoS Pathog 9, e1003124 10.1371/journal.ppat.1003124

40. Rahman, K., Datta, S. A. K., Beaven, A. H., Jolley, A. A., Sodt, A. J., and Compton, A. A. (2022) Cholesterol Binds the Amphipathic Helix of IFITM3 and Regulates Antiviral Activity J Mol Biol 434, 167759 10.1016/j.jmb.2022.167759

41. Hach, J. C., McMichael, T., Chesarino, N. M., and Yount, J. S. (2013) Palmitoylation on conserved and nonconserved cysteines of murine IFITM1 regulates its stability and anti-influenza A virus activity J Virol 87, 9923–9927 10.1128/JVI.00621-13

42. Percher, A., Ramakrishnan, S., Thinon, E., Yuan, X., Yount, J. S., and Hang, H. C. (2016) Mass-tag labeling reveals site-specific and endogenous levels of protein S-fatty acylation Proc Natl Acad Sci U S A 113, 4302–4307 10.1073/pnas.1602244113

43. Yount, J. S., Karssemeijer, R. A., and Hang, H. C. (2012) S-palmitoylation and ubiquitination differentially regulate interferon-induced transmembrane protein 3 (IFITM3)-mediated resistance to influenza virus J Biol Chem 287, 19631–19641 10.1074/jbc.M112.362095

44. Yount, J. S., Moltedo, B., Yang, Y. Y., Charron, G., Moran, T. M., Lopez, C. B. et al. (2010) Palmitoylome profiling reveals S-palmitoylation-dependent antiviral activity of IFITM3 Nat Chem Biol 6, 610–614 10.1038/nchembio.405

45. Rahman, K., Coomer, C. A., Majdoul, S., Ding, S. Y., Padilla-Parra, S., and Compton, A. A. (2020) Homology-guided identification of a conserved motif linking the antiviral functions of IFITM3 to its oligomeric state Elife 9, 10.7554/eLife.58537

46. Wilt, I., Jolley, A. A., Rahman, K., Lai, K. K., Shi, G., Andresson, T., et al. (2025) IFITM1 and IFITM3 cooperate to restrict virus entry in endolysosomes bioRxiv 10.1101/2025.06.01.657267

47. Winkler, M., Wrensch, F., Bosch, P., Knoth, M., Schindler, M., Gartner, S., et al. (2019) Analysis of IFITM-IFITM Interactions by a Flow Cytometry-Based FRET Assay Int J Mol Sci 20, 10.3390/ijms20163859

48. Pelmenschikov, V., Blomberg, M. R., and Siegbahn, P. E. (2002) A theoretical study of the mechanism for peptide hydrolysis by thermolysin J Biol Inorg Chem 7, 284–298 10.1007/s007750100295

49. Karthigeyan, K. P., Zhang, L., Loiselle, D. R., Haystead, T. A. J., Bhat, M., Yount, J. S., et al. (2021) A bioorthogonal chemical reporter for fatty acid synthase-dependent protein acylation J Biol Chem 297, 101272 10.1016/j.jbc.2021.101272

50. Lin, D. T., and Conibear, E. (2015) ABHD17 proteins are novel protein depalmitoylases that regulate N-Ras palmitate turnover and subcellular localization Elife 4, e11306 10.7554/eLife.11306

51. Miki, T., and Orii, Y. (1985) The reaction of horseradish peroxidase with hydroperoxides derived from Triton X-100 Anal Biochem 146, 28–34 10.1016/0003-2697(85)90390-2

52. McKenna, M. J., Adams, B. M., Chu, V., Paulo, J. A., and Shao, S. (2022) ATP13A1 prevents ERAD of folding-competent mislocalized and misoriented proteins Mol Cell 82, 4277–4289 e4210 10.1016/j.molcel.2022.09.035

53. Guna, A., Hazu, M., Pinton Tomaleri, G., and Voorhees, R. M. (2023) A TAle of Two Pathways: Tail-Anchored Protein Insertion at the Endoplasmic Reticulum Cold Spring Harb Perspect Biol 15, 10.1101/cshperspect.a041252

54. Hegde, R. S., and Keenan, R. J. (2024) A unifying model for membrane protein biogenesis Nat Struct Mol Biol 31, 1009–1017 10.1038/s41594-024-01296-5

55. Ji, J., Cui, M. K., Zou, R., Wu, M. Z., Ge, M. X., Li, J. et al. (2024) An ATP13A1-assisted topogenesis pathway for folding multi-spanning membrane proteins Mol Cell 84, 1917–1931 e1915 10.1016/j.molcel.2024.04.010

56. Kamemura, K., Kozono, R., Tando, M., Okumura, M., Koga, D., Kusumi, S., et al. (2024) Secretion of endoplasmic reticulum protein VAPB/ALS8 requires topological inversion Nat Commun 15, 8777 10.1038/s41467-024-53097-5

57. Wang, J., Han, S., and Ye, J. (2023) Topological regulation of a transmembrane protein by luminal-to-cytosolic retrotranslocation of glycosylated sequence Cell Rep 42, 112311 10.1016/j.celrep.2023.112311

58. Dederer, V., and Lemberg, M. K. (2021) Transmembrane dislocases: a second chance for protein targeting Trends Cell Biol 31, 898–911 10.1016/j.tcb.2021.05.007

59. McKenna, M. J., and Shao, S. (2023) The Endoplasmic Reticulum and the Fidelity of Nascent Protein Localization Cold Spring Harb Perspect Biol 15, 10.1101/cshperspect.a041249

60. Tipper, D. J., and Harley, C. A. (2023) Spf1 and Ste24: quality controllers of transmembrane protein topology in the eukaryotic cell Front Cell Dev Biol 11, 1220441 10.3389/fcell.2023.1220441

61. McKenna, M. J., Sim, S. I., Ordureau, A., Wei, L., Harper, J. W., Shao, S., et al. (2020) The endoplasmic reticulum P5A-ATPase is a transmembrane helix dislocase Science 369, 10.1126/science.abc5809

62. Coy-Vergara, J., Rivera-Monroy, J., Urlaub, H., Lenz, C., and Schwappach, B. (2019) A trap mutant reveals the physiological client spectrum of TRC40 J Cell Sci 132, 10.1242/jcs.230094

63. Bhat, N. H., Vass, R. H., Stoddard, P. R., Shin, D. K., and Chien, P. (2013) Identification of ClpP substrates in Caulobacter crescentus reveals a role for regulated proteolysis in bacterial development Mol Microbiol 88, 1083–1092 10.1111/mmi.12241

64. Flynn, J. M., Neher, S. B., Kim, Y. I., Sauer, R. T., and Baker, T. A. (2003) Proteomic discovery of cellular substrates of the ClpXP protease reveals five classes of ClpX-recognition signals Mol Cell 11, 671–683 10.1016/s1097-2765(03)00060-1

65. Westphal, K., Langklotz, S., Thomanek, N., and Narberhaus, F. (2012) A trapping approach reveals novel substrates and physiological functions of the essential protease FtsH in Escherichia coli J Biol Chem 287, 42962–42971 10.1074/jbc.M112.388470

66. Cho, N. H., Cheveralls, K. C., Brunner, A. D., Kim, K., Michaelis, A. C., Raghavan, P., et al. (2022) OpenCell: Endogenous tagging for the cartography of human cellular organization Science 375, eabi6983 10.1126/science.abi6983

67. Chen, Y. J., Knupp, J., Arunagiri, A., Haataja, L., Arvan, P., and Tsai, B. (2021) PGRMC1 acts as a size-selective cargo receptor to drive ER-phagic clearance of mutant prohormones Nat Commun 12, 5991 10.1038/s41467-021-26225-8

68. Kim, J. Y., Kim, S. Y., Choi, H. S., An, S., and Ryu, C. J. (2019) Epitope mapping of anti-PGRMC1 antibodies reveals the non-conventional membrane topology of PGRMC1 on the cell surface Sci Rep 9, 653 10.1038/s41598-018-37441-6

69. McGuire, M. R., and Espenshade, P. J. (2023) PGRMC1: An enigmatic heme-binding protein Pharmacol Ther 241, 108326 10.1016/j.pharmthera.2022.108326

70. Jimenez-Munguia, I., Beaven, A. H., Blank, P. S., Sodt, A. J., and Zimmerberg, J. (2022) Interferon-induced transmembrane protein 3 (IFITM3) and its antiviral activity Curr Opin Struct Biol 77, 102467 10.1016/j.sbi.2022.102467

71. Xie, Q., Wang, L., Liao, X., Huang, B., Luo, C., Liao, G. et al. (2024) Research Progress into the Biological Functions of IFITM3 Viruses 16, 10.3390/v16101543

